# Innate Immune Activation and Mitochondrial ROS Invoke Persistent Cardiac Conduction System Dysfunction after COVID-19

**DOI:** 10.1101/2024.01.05.574280

**Authors:** Deepthi Ashok, Ting Liu, Joseph Criscione, Meghana Prakash, Byunggik Kim, Julian Chow, Morgan Craney, Kyriakos N. Papanicolaou, Agnieszka Sidor, D. Brian Foster, Andrew Pekosz, Jason Villano, Deok-Ho Kim, Brian O’Rourke

## Abstract

**Background:** Cardiac risk rises during acute SARS-CoV-2 infection and in long COVID syndrome in humans, but the mechanisms behind COVID-19-linked arrhythmias are unknown. This study explores the acute and long term effects of SARS-CoV-2 on the cardiac conduction system (CCS) in a hamster model of COVID-19.

**Methods:** Radiotelemetry in conscious animals was used to non-invasively record electrocardiograms and subpleural pressures after intranasal SARS-CoV-2 infection. Cardiac cytokines, interferon-stimulated gene expression, and macrophage infiltration of the CCS, were assessed at 4 days and 4 weeks post-infection. A double-stranded RNA mimetic, polyinosinic:polycytidylic acid (PIC), was used in vivo and in vitro to activate viral pattern recognition receptors in the absence of SARS-CoV-2 infection.

**Results:** COVID-19 induced pronounced tachypnea and severe cardiac conduction system (CCS) dysfunction, spanning from bradycardia to persistent atrioventricular block, although no viral protein expression was detected in the heart. Arrhythmias developed rapidly, partially reversed, and then redeveloped after the pulmonary infection was resolved, indicating persistent CCS injury. Increased cardiac cytokines, interferon-stimulated gene expression, and macrophage remodeling in the CCS accompanied the electrophysiological abnormalities. Interestingly, the arrhythmia phenotype was reproduced by cardiac injection of PIC in the absence of virus, indicating that innate immune activation was sufficient to drive the response. PIC also strongly induced cytokine secretion and robust interferon signaling in hearts, human iPSC-derived cardiomyocytes (hiPSC-CMs), and engineered heart tissues, accompanied by alterations in electrical and Ca^2+^ handling properties. Importantly, the pulmonary and cardiac effects of COVID-19 were blunted by in vivo inhibition of JAK/STAT signaling or by a mitochondrially-targeted antioxidant.

**Conclusions:** The findings indicate that long term dysfunction and immune cell remodeling of the CCS is induced by COVID-19, arising indirectly from oxidative stress and excessive activation of cardiac innate immune responses during infection, with implications for long COVID Syndrome.

## Introduction

With over 1 million confirmed deaths in the US and almost 7 million globally, the COVID-19 pandemic uncovered a significant gap in our understanding of virus-host interactions and how to prevent the systemic complications of infection. Much is known about the array of mechanisms involved in antiviral innate immune responses in general, and how some viruses evade these defenses. However, the specific mechanisms underlying the short- and long-term cardiac consequences of RNA viruses like SARS-CoV-2 are not well understood. Adverse cardiac electrophysiological effects are a prominent feature of SARS-CoV infection(*1, 2*), including changes in QT interval(*3*), atrial arrhythmias(*1, 4, 5*), bradyarrhythmias, ventricular tachycardias (VT), and cardiac arrest(*6, 7*). A Heart Rhythm Society survey found that atrial fibrillation was the most common tachyarrhythmia (21%), with severe sinus bradycardia (8%) and complete heart block (6%) as the most common bradyarrhythmia. Ventricular arrhythmias included premature ventricular complexes (5.3%), non-sustained VT (6.3%), monomorphic VT (3.8%), polymorphic VT (Torsade de Pointes, 3.5%), or cardiac arrest with VT/VF (4.8%)(*8*). Thus, cardiovascular complications may be an important contributor to SARS-CoV-2 morbidity, even extending to time points well after the initial infection is resolved (long COVID syndrome). Interestingly, cardiac inflammation sometimes develops or persists after recovery from COVID-19, as reported for collegiate athletes post-infection, even among asymptomatic subjects(*9*).

The mechanisms underlying these COVID-19-associated arrhythmias are unknown and could be due to either direct or indirect effects of the virus. For example, during the SARS-CoV outbreak in Toronto in 2002, up to 35% of human heart autopsy samples were positive for SARS-CoV RNA and showed evidence of macrophage infiltration and myocyte necrosis(*10*). Similarly, several early studies reported SARS-CoV-2 RNA present in post-mortem myocardial tissue from COVID-19 patients(*11, 12*), although it is unclear whether this represents viral infection or RNA present in interstitial infiltrates from the blood, secondary to vascular leakage(*13*). A significant fraction (up to 60%)(*14*) of SARS-CoV-2 infected patients showed evidence of cardiac inflammation or injury, with evidence of increased plasma NT-ProBNP, Troponin T, and IL-6(*15–19*). Nevertheless, the overall incidence of fulminant myocarditis during or after COVID-19 infection is likely to be low(*20*), and more recent studies do not support widespread viral replication in the heart(*21–24*). Even in the absence of direct infection, cardiac interferon-stimulated gene (ISG) transcripts, including 2′-5′-Oligoadenylate (2-5A) Synthetase (OAS), significantly increase, indicating that antiviral innate immunity is induced, along with marked changes in mitochondrially-encoded genes(*22*). These data support the idea that SARS-CoV-2 can affect the heart by triggering an organism-wide antiviral response, but whether this is protective or detrimental (i.e., the “cytokine storm” model) is difficult to ascertain. Outcomes may depend on timing, as early activation of innate immunity inhibits viral replication, but sustained activation might impair function(*25*).

Oxidative stress is an important component of inflammatory and immune cell responses, and excess reactive oxygen species (ROS) or markers of oxidative stress have been documented for a number of respiratory viral infections, including COVID-19(*26*). Human studies have associated serum antioxidant depletion with COVID-19 severity(*27, 28*), suggesting that antioxidant interventions might prevent or reverse the pathogenesis. Several clinical trials were implemented to assess the benefits of antioxidant therapy, including nonenzymatic dietary antioxidants (vitamins A, E, C, Zinc, and selenium) (*29*), reduced glutathione (GSH) or its precursor N-Acetylcysteine (NAC)(*30, 31*), nitric oxide (NCT04388683), natural products(*32*), or synthetic antioxidants (Tempol; NCT04729595). These antioxidant interventions typically decreased cytokines or biomarkers of COVID-19 but had little effect on clinical outcomes. None of the studies were designed to assess effects of the antioxidant treatments on pulmonary or cardiac function in detail, or their long-term impact.

Here, we examine the acute and persistent cardiac electrophysiological effects after intranasal SARS-CoV-2 infection in hamsters, while assessing pulmonary function in parallel during the course of COVID-19. Even in the absence of detectable viral protein expression in the heart, we find pronounced expression of interferon-stimulated genes (ISG), cardiac arrhythmias, and immune cell remodeling in the cardiac conduction system (CCS). Interestingly, similar arrhythmias could be induced by a mimetic of the viral double-stranded RNA (dsRNA)-triggered innate immune response in naïve animals. Guided by in vitro studies of innate immune activation in human iPSC-derived cardiomyocyte monolayers, we further show that in vivo interventions designed to inhibit interferon signaling or mitochondrial ROS mitigate the cardiac and pulmonary effects of COVID-19 in the hamster model.

## Materials and Methods

### Study design

The objective of this study was to mechanistically investigate COVID-19 induced arrhythmias. For this study, we chose a previously established Syrian golden hamster model of SARS-CoV-2 that mirrors the pulmonary tissue damage and inflammation phenotype as seen in humans(*36*). Male hamsters 8-10 weeks of age were included in this study. Sample size was determined by space availability in BSL-3 labs as well as previously established studies in hamsters(*106–108*). In general, 3-11 hamsters per treatment group per parameter were included in this study. Only one hamster that had an early-death due to trauma induced after telemetry-implant surgery was excluded from this study. We included data points from all the other hamsters. Hamsters were divided into four groups: mock treated, intranasal SARS-CoV-2 infected, SARS-CoV-2 infected with Ruxolitinib administration and SARS-CoV-2 infected with mitoTEMPO administration. Further, we wanted to test our hypothesis that arrhythmias were caused by activation of innate immune response in the heart in the absence of viral infection. Guinea pigs were used in this section of the study since they closely match the electrophysiological characteristics of human heart. 3 male guinea pigs used as the control group and 7 male guinea pigs were injected with PIC. Some outliers in qPCR experiments were excluded based on if they were above or below the interquartile range (IQR). We used data only between the first quartile (Q1) and third quartile (Q3). Any datapoint below Q1-1.5*IQR or above Q3+1.5*IQR were excluded. As for hiPSC-CM studies, data were collected from 3 different batches of differentiated cells.

### Animal models

The animal procedures in this study were approved by the Institutional Animal Care and Use Committee of Johns Hopkins University, in accordance with the National Research Council’s Guide for the Care and Use of Laboratory Animals (eighth edition). The Johns Hopkins animal care and use program is accredited by the Association for Assessment and Accreditation of Laboratory Animal Care International.

SARS-CoV-2 infection experiments with Golden Syrian hamsters were performed in a biosafety level 3 facility at Johns Hopkins Research Animal Resources, in compliance with the established ethical guidelines.

2.4X10^7^ TCID50(50% tissue culture infectious dose)/ml of SARS-CoV-2/Delta variant (SARS-CoV-2/USA/MD-HP05660/2021; GIAID accession number EPI_ISL_2331507) in 100 µl Dulbecco’s modified Eagle medium (DMEM) was intranasally administered (50 µl per nare) to male hamsters (Envigo, Haslett MI) as described previously (*36*). Mock control animals received DMEM alone.

Cardiac injection of PIC in guinea pigs was performed as described previously(*109*). Briefly, 450-500g male Hartley guinea pigs (HillTop Lab Animals, PA) were anesthetized with 4% isoflurane for 4 min, intubated, and ventilated with oxygen and 2% isoflurane for direct myocardial injection. 200 µl PBS with 100 µg PIC was intramuscularly injected into the ventricular wall at multiple sites. Mock control animals received PBS only. For studies of isolated cardiomyocytes in Figure 5J, a 100 µl solution containing 1×10^10^ PFU adenoviral vector expressing GFP as an indicator, with or without 50 µg PIC, was injected into the ventricular wall around the apex. Cardiomyocytes were isolated as described previously (*109*), at 3 days after injection. Physiological studies were performed with GFP-positive cells.

### Radiotelemetry device and osmotic pump implantation

Device implantation in hamsters was performed 4 days before the SARS-CoV-2 infection. 6-to 8-week-old male hamsters were anesthetized with 4% isoflurane in a closed chamber for 4 min and then maintained with oxygen and 2% isoflurane during surgery. To measure subpleural pressure, the esophagus was isolated approximately 1.5 cm from its junction with the diaphragm and a 25-gauge needle was inserted between the serosal and muscularis layers. The needle was tunneled past the junction with the diaphragm into the thoracic cavity. The needle was then removed and the transmitter catheter of the dual pressure/biopotential device (model HD-X11, DSI, St. Paul MN) was threaded through the tunnel and secured in place with suture on the serosal layer. To monitor the ECG, the biopotential leads (Model: HD-X11 or ETA-F10, DSI, St. Paul MN) were secured with suture on the muscle layer at Lead II position. The body of the device was placed on the side of abdominal cavity. For treatment with mitoTEMPO or Ruxolitinib (Ruxo), Alzet osmotic pumps filled with drug or vehicle were implanted into abdominal cavity to deliver the drug for 14 days. Both mitoTEMPO and Ruxo were delivered at 1 mg/kg/day. After 3 days of recovery, baseline ECG and subpleural pressure were recorded for 24 hr. Then the hamsters were sedated intramuscularly with xylazine and ketamine and inoculated intranasally with 105 50% TCID50 of SARS-CoV-2 delta strain diluted in 100 μL Dulbecco’s modified Eagle’s medium. Uninfected animals were mock inoculated with 100 μL of Dulbecco’s modified Eagle’s medium intranasally to serve as controls. ECG was recorded for 24 hr at 1, 3, 5, 7, 14, 21, and 28 days post inoculation (DPI). Hamsters were euthanized at 4 or 28 DPI. Body weights were measured daily until 10 DPI, then on 14, 21, and 28 DPI.

Implantation of the biopotential devices (Model ETA-F10, DSI, St. Paul MN) in guinea pigs was performed in the same surgical session as the PIC injection. ECGs were recorded at 2 and 3 days after injection and the animals were then euthanized for heart tissue collection.

### Radiotelemetry data analysis

ECG and subpleural pressure data were collected with Ponemah 3.0 (DSI, St. Paul MN). AV block was manually counted in 2-hour duration from 0-2am. Mean RR interval and frequency of RR interval longer than meanRR+100ms for hamster or meanRR+2xS.D. for guinea pig was analyzed with custom-written Matlab routines (Mathworks, Inc.). Subpleural pressure and HRV were analyzed with LabChart 8 (AD Instruments, Colorado Springs CO) with Modules of Pressure and HRV.

### Cultured cell models

Human induced pluripotent stem cell derived cardiomyocytes (hiPSC-CMs) were differentiated according to previously established protocols (*110*). The hiPSC-CMs were then enriched to >95% purity using Miltenyi’s MACS purification protocol. Briefly, hiPSC-CMs on day 10-14 of differentiation were trypsinized and collected in a tube. Dead cells were removed using a dead cell removal kit (Catalog No. 130-090-101, Miltenyi Biotec). Cardiomyocytes were purified using a PSC-Derived Cardiomyocyte Isolation Kit (130-110-188, Miltenyi Biotec). LS columns were used to enrich cardiomyocytes (130-042-401, Miltenyi Biotec). Purified cardiomyocytes were then plated in monolayers onto Geltrex™ coated (Cat. No. A1413202, ThermoFisher Scientific) plates for further experiments. A549 lung carcinoma epithelial cells were used to study effects of innate immune activation in a human lung-derived cell model.

### Polyinosinic–polycytidylic acid activation of innate immune signaling

Polyinosinic–polycytidylic acid (PIC) is a dsRNA mimetic known to activate the intracellular pattern recognition receptors OAS/MDA5/RIG-I to trigger IRF3-mediated signaling, leading to the upregulation of Type I Interferons. PIC (Cat. No. P1530, Sigma) was reconstituted in saline at 10mg/ml or 20mg/ml and used at 200μg/ml in our cellular experiments. In A549 cells, PIC treatment alone elicited a weak interferon response (indexed by STAT1, pSTAT1, OAS1,2,3 protein expression) that was markedly accentuated by co-addition of an empty replication-incompetent adenovirus vector (AdV; Supplemental Fig. 1). This effect was likely due to enhanced AdV-induced endocytosis(*111*) and internalization of PIC, as it could be mimicked by lipofectamine treatment. AdV alone did not activate the immune response and was not cytotoxic (unlike lipofectamine), so subsequent immune challenges were performed with the combination of PIC+AdV in A549 cells. However, in hiPSC-CMs, PIC alone was enough to mount a substantial immune response with STAT1 and pSTAT1 increase (Supplemental figure 2). Addition of AdV, did not significantly increase immune response. Therefore, the experiments in cardiomyocytes were performed with PIC alone, unless specifically noted in the figures (e.g., Figures 5D, E and J), when it was co-applied with AdV.

### Protein extraction and Immunoblotting

Ventricles and lungs were harvested at 4 or 28 DPI. Tissues were rinsed in cold PBS, rapidly heat-stabilized (Stabilizor^TM^, Denator, Inc.), snap-frozen in liquid nitrogen, and stored in a -80 freezer. To extract protein, stabilized tissues were homogenized with RIPA buffer in the presence of 2% SDS, solubilized, and boiled in 1x LDS sample buffer for SDS-PAGE. The protein mixture was separated on a 4-12% NuPAGE gel (1 mm, Invitrogen). Samples were run at room temperature for 35 min at 200 V. Proteins were transferred to nitrocellulose membranes with iBlot (Invitrogen, Inc.), using program 3 for 7 min. Membranes were stained with Ponceau S solution (Sigma-Aldrich) to evaluate the transfer efficiency. Membranes were blocked for 1 h using Odyssey® blocking buffer (Li-Cor Biosciences) and incubated with primary antibody (Supplementary Table 1) overnight at 4°C. Antibody binding was visualized with an infrared imaging system using IRDye secondary antibodies (800CW Donkey anti-Rabbit IgG, #926-32211 and 680RD Donkey anti-Mouse IgG, #926-68072; Odyssey, Licor Biosciences) and quantification of band intensity was performed using the Odyssey Application Software 3.0.

hiPSC-CMs, A549 cell lysates or EHTs were collected 96 hours after treatment with PIC (200µg/ml) or PIC-AdV or PIC+Ruxolitinib (1μM) or PIC+mitoTempo (1μM) along with an untreated control group. Four days after treatment, cells were washed in ice-cold PBS and lysed in RIPA buffer (Cat. No. R0278, Sigma) supplemented with protease (Cat. No. P8340 or Cat. No. 11836170001 Sigma) and phosphatase (Cat. No. P0044 or Cat. No. 4906845001, Sigma) inhibitor cocktail. Engineered Heart Tissues were snap-frozen in liquid nitrogen and stored in the -80 freezer. Lysates were sonicated and centrifuged to spin down the insoluble pellet. Supernatants were collected to perform immunoblotting. Protein samples were denatured at 70° C for 10 min after combining with 5% b-mercaptoethanol and 1X NuPAGE™ LDS Sample Buffer (Cat. No. NP0007, ThermoFisher) and run in 4-12% Bis-Tris gels (Cat. No. WG1402BOX or NP0322BOX). Proteins were transferred to nitrocellulose membranes using iBlot transfer stacks (Cat. No. IB301001). Membranes were blocked with Intercept® TBS blocking buffer (Cat. No. 927-60001, LICOR) for 1 hour at room temperature. Membranes were incubated with primary antibodies (diluted 1:1000 in blocking buffer) overnight at 4°C. A list of primary antibodies used is provided in Supplemental Table 1. Membranes were incubated with secondary antibodies (IR Dye 800w goat anti-rabbit, Cat. No. 926-32211, LI-COR) at 1:10000 dilution in Intercept ® blocking buffer for 1 hour at room temperature. For detection, we used the LICOR-Odyssey system. Normalization for protein loading was based on intensity staining of the membrane with Ponceau. Band intensities were analyzed and quantified using NIH-ImageJ software (*112, 113*)

### Immunofluorescence

Tissues were fixed with 4% paraformaldehyde for 3 days and transferred to 30% sucrose for 24-48 hours at 4°C, followed by OCT embedding. For immunofluorescence, frozen sections were washed with PBS, blocked with 2% BSA and 0.05% triton X-100 in PBS for 1 hr, and incubated with primary antibodies at 4°C overnight and secondary antibodies for 1 hr. Images were acquired with a spinning-disk confocal microscope (Andor Revolution) and analyzed with ImageJ. Cell numbers were normalized to tissue area.

### qPCR

Total RNA was isolated from heat-stabilized tissue using Trizol regent following manufacture’s manual. Equal amounts of RNA were transcribed into cDNA by High-Capacity cDNA Reverse Transcription Kit (Applied Biosystems). cDNA product was 1 to 40 diluted and real-time PCR was performed using FastStart SYBR Green Master Mix (Roche) on QuantStudio 5 (ThermoFisher) with primers specific to hamster genes encoding CCL2, CXCL10, CXCL11, IFNα, IFNβ, IFNγ, IL1b, IL6, IL10, OAS1, OAS2, OAS3, RIG1, TGFβ, TNFα as well as β-actin as an internal reference (sequence of primers in attached spreadsheet, Supplemental Table 2).

### Cytokine Proteome Profiler

Cell-culture supernatants were collected 96 hr after PIC or PIC+ Ruxolitinib, or PIC+ mitoTEMPO treatment. Cytokines released from hiPSC-CMs into the supernatants were analyzed using Proteome Profiler Human XL Cytokine Array Kit (Cat. No. ARY022B R&D Systems). Chemiluminescence from the membrane was detected and scanned on an iBright Imaging System FL1000 (Thermo Fisher Scientific). Scanned images were analyzed using Quick Spots image analysis software, version 25.5.2.3 (Ideal Eyes Systems).

### Ca^2+^ transient measurements

250,000 hiPSC-CMs were plated per well onto 8-well plastic plates (ibidi, Inc. Cat. No. 80806). A working stock solution of 2mM Fluo-4AM (Catalog No. F14201, Invitrogen) was made by dissolving in equal parts DMSO and 20% (w/v) Pluronic® F-127 (Catalog No. P3000MP, Invitrogen). hiPSC-CMs were loaded with 2μM Fluo-4 AM in cell culture media and incubated at 37°C for 15 minutes. Cells were then washed once with PBS before finally incubating them in Tyrode’s buffer (130 mM NaCl, 5 mM KCl, 1 mM MgCl2, 10 mM NaHEPES, 1 mM CaCl2 and 5 mM Glucose) for imaging. Ca^2+^ transients from hiPSC-CMs were imaged on a spinning disk-confocal microscope (Andor Revolution, Olympus IX-70). Images were collected for 25 seconds. Ca^2+^ transients were monitored at 20X magnification with an Olympus UCPlanFL N objective. 488nm laser excitation was used at 35% power, 30msec exposure and 33.3 frames per second over a duration of 22-25 seconds. Custom-written ImageJ and excel-based macros were used to analyze Ca^2+^ transient dynamics.

### Engineered Heart Tissues

Engineered heart tissues (EHTs) were fabricated as per previously described protocols(*114*). Briefly, a mixture of 1×10^6^ hiPSC-CMs (Celo.Cardiomyocytes, Celogics, Washington,USA), 5×10^4^ HS-27A human bone marrow stromal cells (ATCC), and 0.5mg human fibrinogen (Sigma-Aldrich) in a total volume of 100µL EHT culture media (Celo.Cardiomyocyte advanced culture media, B27, 5g/L aminocaproic acid) with 10µM ROCKi, was prepared per tissue. This mixture was then pipetted into Mantarray casting wells containing 50µL EHT culture media with 0.3U human thrombin (Sigma-Aldrich). EHTs were then incubated for one hour at 37°C, at which point 1mL of EHT culture media with 10µM ROCKi was added to each casting well. After overnight incubation at 37°C EHTs were transferred to 2mL/well EHT culture media. EHT culture media was changed every 2-3 days thereafter.

### Sarcomere shortening and Ca^2+^ transient recordings of adult cardiomyocytes

Ventricular myocytes were loaded with 3 µM Fura2-AM (Invitrogen, Molecular Probes, Carlsbad CA) in a modified Tyrode’s solution containing (in mM) NaCl 138, KCl 4, CaCl_2_ 2, MgCl_2_ 1, HEPES 10, NaH_2_PO_4_ 0.33, and Glucose 10 (pH 7.4 with NaOH) for 15min. After rinsing, cells were placed in a perfusion chamber with a flow-through rate of 2 ml/min, and sarcomere length and whole cell Ca^2+^ transients were recorded using an inverted fluorescence microscope (Nikon, TE2000), and IonOptix (Myocam®) software.

### Statistical analysis

Values are expressed as mean ± SEM. Statistical analyses between 2 groups or among multiple groups were performed with Student’s t-test or 1-way ANOVA using GraphPad Prism version 10.0.2 for MacOSX. The time series data from the hamster studies were analyzed with parametric (Figure 1, Figure 2B, Figure 7 A-D) or non-parametric methods (Figure 2C-E and Figure 7 E-G) depending on the normality of the data distribution. Parametric analysis was performed using JMP Pro 16.0. The responses of body temperature, respiration, and RR intervals to SARS-CoV2 infection, with or without treatment, within each group were analyzed with 1-way ANOVA with repeated measures followed by Dunnet’s test to compare each time point with baseline (D0). The comparisons among groups were analyzed with multiple regression followed by Tukey HSD test for pairwise comparison. Non-parametric analysis was performed using R. The time course response within each group was analyzed with Friedman’s test and the comparison among groups was analyzed with the Kruskal-Wallis test. Conover test was used for pairwise comparison in both methods. ECG signals recorded for 24 hours for each time point were analyzed by 6-hour segment. RR interval was analyzed with a nested model of multiple regression. In non-parametric analysis, the parameters of the 6-hr segments were averaged for each time point. Sample size is provided in figure legends. A complete list of results of statistical analysis and p-value for all the parameters for different hamster groups is presented in supplementary table 4 (S4).

**Fig. 1.**
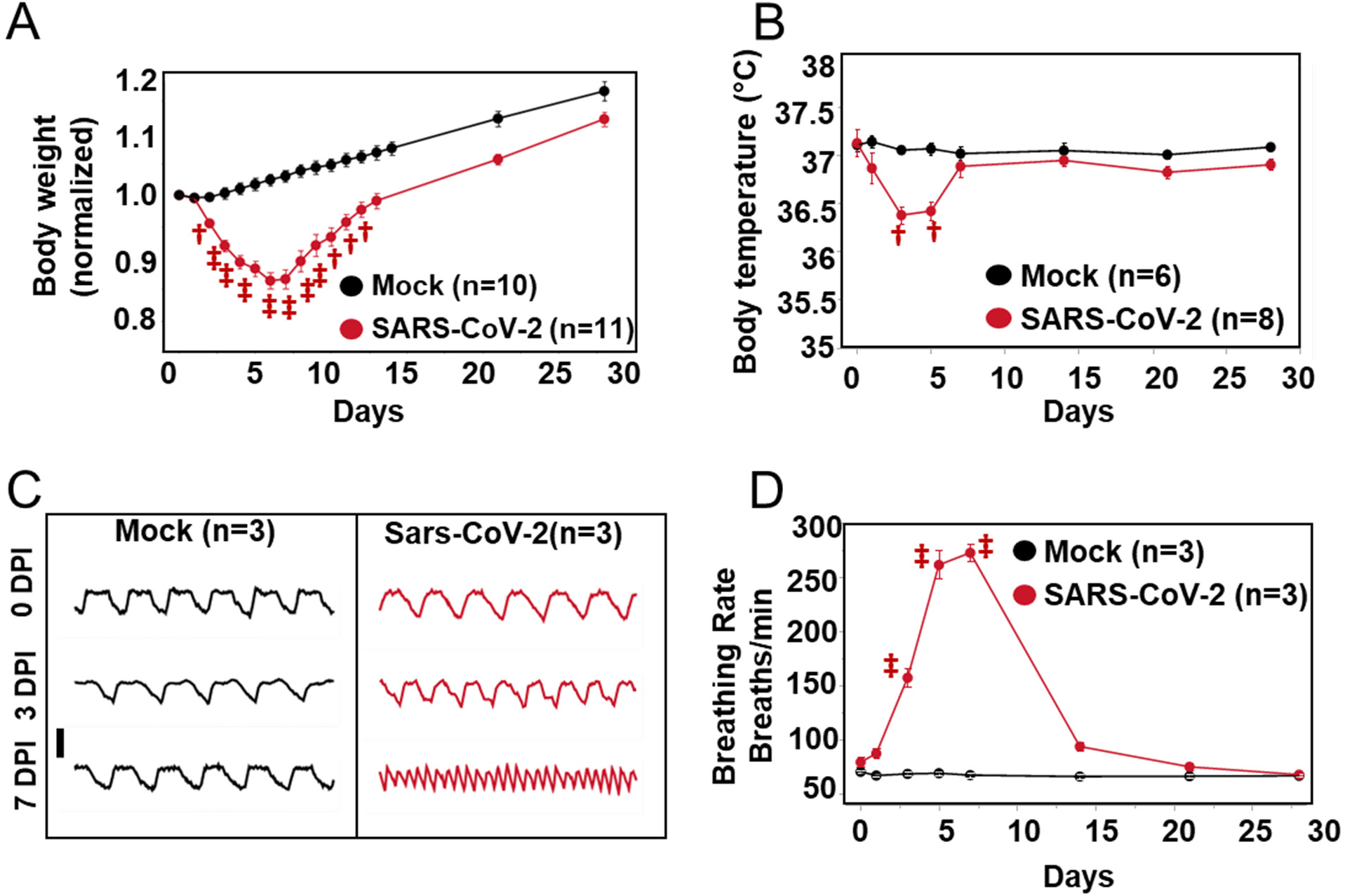
SARS-CoV-2 infection causes weight loss, hypothermia, and tachypnea in the hamster model. **(A)** Body weight decreased after infection, reached a minimum at 6-7 dpi, and then recovered back to normal. **(B)** SARS-CoV-2 Infection induced a transient decrease in body temperature peaking at 3 dpi and recovering by 7 dpi. **(C)** Representative traces of subpleural pressure recorded in Mock-infected (left) and SARS-CoV-2 (right)-infected hamsters at 0 (upper), 3 (middle), and 7 (lower) dpi. Vertical bar: 10mmHg; horizontal bar: 1 sec. **(D)** Breathing rate increased following SARS-CoV-2 infection, peaked at 7 dpi, and recovered by 14 dpi. ‡ p<0.0001

**Fig. 2.**
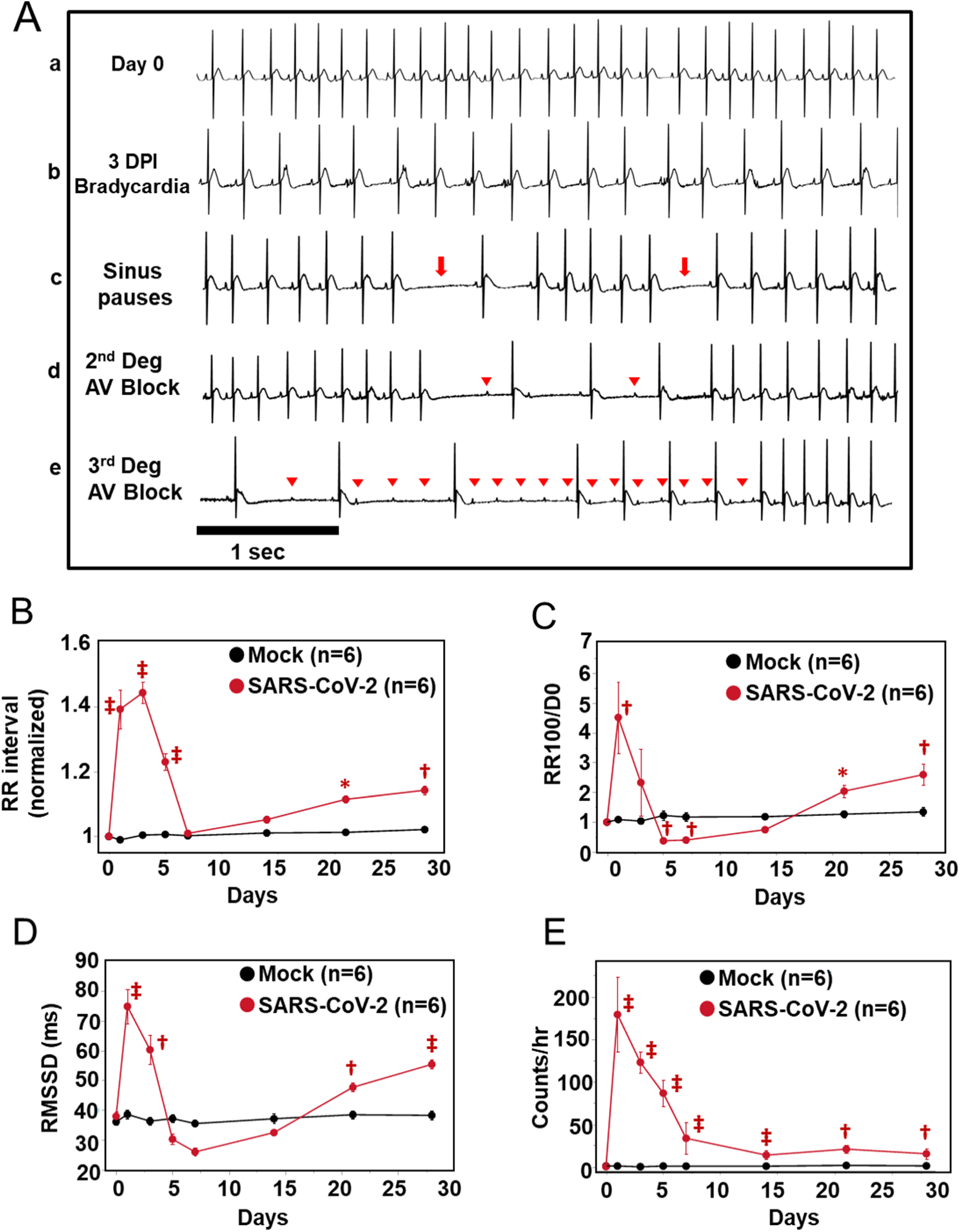
Effects of SARS-CoV-2 infection on cardiac electrophysiology in the hamster model. **(A)** Representative ECG recording showing normal baseline rhythm (a) and arrhythmias induced by SARS-CoV-2 infection including bradycardia (b), sinus pauses (c, arrows), 2^nd^ degree AV block (d) and 3^rd^ degree AV block (e). Arrowheads indicate p waves. Horizontal bar: 1 sec. **(B-E)** Analysis of ECGs from 0 to 28 dpi typically showed a triphasic pattern of SARS-CoV-2 effects on cardiac rhythm: an acute peak within 7 dpi, recovery to, or below, baseline, and a long-term effect developing between 7 dpi and 28 dpi. Mean RR interval (normalized to D0) **(B)**, incidence of long sinus pauses (RR>meanRR+100ms; normalized to D0) **(C)**, and RMSSD **(D)** peaked at 1-3 dpi, and returned to levels close to or lower than baseline, and then gradually increased to levels significantly higher than baseline at 21 and 28 dpi. The rate of AV Block events **(E)** peaked early and did not return to baseline level, remaining significantly higher than baseline between 7 and 28 dpi. Red symbols denoting p-values * p<0.05; † p<0.01; ‡ p<0.0001 on the figures compare mock vs. SARS-CoV2 infected groups. P-values comparing SARS-CoV-2 data from day 0 through day 28 are available in Supplementary table 4.

## Results

### Effects of SARS-CoV-2 infection on systemic and pulmonary function in hamsters

In humans, systemic inflammatory responses to pathogen infections often involve either hyper- or hypothermia, tachypnea, and weight loss (*33*). While the majority (∼80%) of COVID-19 patients present with early fever, hypothermia(*34*) and weight loss(*35*) are also associated with poor prognosis. In the hamster COVID-19 model, intranasal inoculation with SARS-CoV-2 (Delta strain) resulted in weight loss, tachypnea and hypothermia within the first 7 days post-infection (dpi). Consistent with a previous study from our institution (*36*), weight loss was maximal at 6 dpi, with a 14±1% decrease compared to baseline, followed by reversion to a normal rate of growth by 28 dpi compared to the mock-infected group (Fig. 1A).

Dual biopotential/pressure radiotelemetry devices (DataSciences International, MN) were used to monitor the electrocardiogram, subpleural pressure, and body temperature simultaneously in freely moving animals. Body temperature decreased after infection, reaching a minimum at 3-5 dpi (37.1±0.1°C at baseline vs. 36.4±0.1°C at 3 dpi), and then recovered to 37.0±0.1°C by 7 dpi (Fig. 1B). Tachypnea developed rapidly between 1 and 5 dpi, from 81±6 breaths/min at baseline to 276±18 and 272±11 breaths/min at 5 and 7 dpi, respectively, and then recovered to 101±3 breaths/min by day 14 (Fig.1 C, D).

### Triphasic effects of COVID-19 infection on cardiac arrhythmias

ECG analysis revealed that SARS-CoV-2 infection resulted in multiple types of cardiac arrhythmias linked to CCS dysfunction, including bradycardia, sinus pauses, and 2nd and 3rd degree atrioventricular (AV) block (Fig.2A). The effects had a triphasic pattern: an early peak at 1-3 dpi, a recovery phase by 7 dpi, and arrhythmia redevelopment that persisted through 28 dpi. As early as 1 day after SARS-CoV-2 infection, marked bradycardia was observed. Compared to baseline, RR interval increased by 39±6% at 1 dpi and peaked at 3 dpi, with an increase of 44±3%, which then reverted to near baseline levels by 7 dpi (Fig. 2B). This was followed by a linear increase in RR interval extending to 4 weeks post-infection that was statistically significant after 21 dpi (11±1%, p<0.001 at 21 dpi and 14±1%, p<0.0001 at 28 dpi) (Fig. 2B). The incidence of beat-to-beat pauses longer than the RR interval mean + 100 msec (RR100) spiked early at 1 dpi (4.5-fold higher than 0 dpi, p<0.0001), undershot the baseline (63% and 60% lower than 0 dpi at 5 and 7 dpi, respectively), and then increased above baseline at 28 dpi (2.6-fold higher than baseline) (Fig. 2C).

Evidence of autonomic nervous system dysfunction was also observed. Heart rate variability analysis revealed that the root mean square of the successive RR differences (RMSSD) was significantly higher than baseline at 1 and 3 dpi and lower at 5 and 7 dpi (74.9 ± 5.7ms at 1 dpi; 60.3 ± 4.9ms at 3 dpi; 30.4±1.7ms at 5 dpi; 26.1 ± 1.4ms at 7 dpi vs. 38.1 ± 1.6ms at 0 dpi), suggesting increased parasympathetic activity followed by impaired vagal responses during the acute COVID phase. RMSSD then increased to levels significantly higher than baseline at 21 and 28 dpi (Fig. 2D). AV block frequency peaked at 1 dpi (179.50±44.53 vs 0.19±0.09 at 0 dpi, p<0.0001) and gradually decreased to 33.56±18.54 (p<0.0001 compared to 0 dpi) at 7 dpi but remained elevated through 28 dpi (Fig. 2E).

### Immune cell remodeling in the cardiac conduction system

Previous reports highlighted the important role of co-localized macrophages in the modulation of specialized pacemaker and conduction system cells in the heart. Resident macrophages were shown to facilitate cardiac conduction(*37*) and enable mitochondrial quality control(*38*), while recruited macrophages contribute to inflammation and cardiac arrhythmias(*39*). Hence, we examined whether immune cell remodeling took place in hearts during COVID-19. Tissue slices encompassing conduction system structures at the atrioventricular septum (at the level of the His bundle) were positive for the conduction system marker Contactin-2 (Cntn2) (Fig. 3A) (*40, 41*). The total number of macrophages (Iba1+), and those expressing (CD163+), a marker of anti-inflammatory or resident macrophages(*42*), were counted and normalized to tissue pixel area in the AV node/His bundle region at both the acute phase (4 dpi) and post-infection phase (28 dpi) (Fig 3B-D). In the mock-infected control heart, the relative densities of Iba-1+ and CD163+ macrophages were higher in the AV/bundle region compared to ventricles (Iba-1+ macrophages: 0.37±0.01 in bundle vs 0.09±0.01 in ventricles, p<0.0001; CD163+ macrophages: 0.16±0.02 in bundle vs 0.03±0.01 in ventricles, p<0.005, see supplement Fig.3). Following SARS-CoV-2 infection at 4 dpi, significant macrophage infiltration was observed throughout the heart and the density of Iba1+ macrophages increased by 78% in the AV/bundle region of COVID-19 animals compared to mock-infected controls (Fig. 3C). Despite the increase in macrophage infiltration, the density of CD163+ macrophages decreased by 74% in the AV/bundle region. After the acute phase, at 28 dpi, Iba1+ macrophage density declined to levels not significantly different from controls (Fig. 3C). CD163+ macrophage density was also not significantly different from controls at 28 dpi, in part because of a decrease in the mock-infected group (Fig. 3D).

**Fig. 3.**
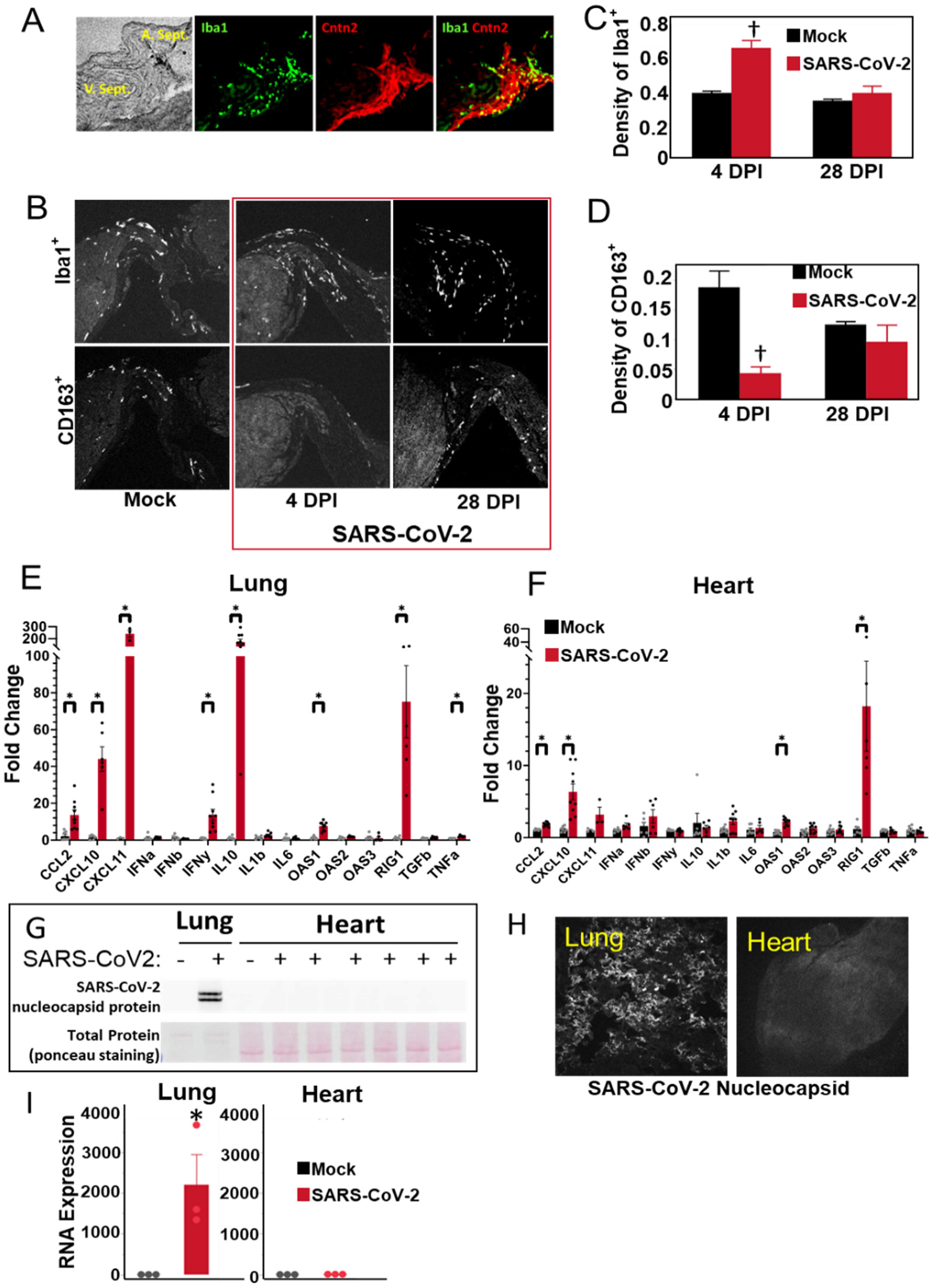
Macrophage remodeling in the cardiac conduction system. **(A)** Representative images showing the conduction tissue marker Contactin-2 (red) and the distribution of Iba-1^+^ macrophages in the region. **(B)** Representative images showing Iba-1^+^ and CD163^+^ macrophages in the conduction region in Mock-infected or SARS-CoV-2-infected hearts at 4 and 28 dpi. **(C-D)** Density of Iba-1^+^ cells and CD163^+^ cells in cardiac conduction system region of the Mock- or SARS-CoV-2-infected hearts at 4 and 28 dpi. † p<0.01. **(E-F)** Gene expression of cytokines and interferon-stimulated genes in lung **(E)** and heart **(F)** at 4 dpi, evaluated with qPCR. * p<0.05; † p<0.005; ‡ p<0.0001. **(G-I)** SARS-CoV-2 nucleocapsid protein was detected in lung, but not in the heart, by western blot (G) or by immunofluorescence **(H)** at 4 dpi. Nucleocapsid mRNA was also detected in lung, but not in the heart, by qPCR **(I)** at 4 dpi. * p<0.05.

### Interferon-stimulated gene and cytokine expression in lung and heart

RT-PCR was used to characterize cytokine expression the lung and the heart. In the lung at 4 dpi, we observed pronounced increases in the anti-inflammatory cytokine IL-10 (150-fold), the monocyte chemoattractant CCL2 (12-fold), and also in IFNγ (12-fold) and downstream interferon-stimulated genes (ISG), including the immune cell chemoattractants CXCL11 (241-fold) and CXCL10 (45-fold) (Fig. 3E), previously linked to poor outcomes in COVID patients (*43–46*). Antiviral innate immune response dsRNA pattern recognition receptors, OAS1 (8-fold) and RIG-I (75-fold), were also elevated in the lung (Fig. 3E). Despite the lack of evidence of widespread myocardial SARS-CoV-2 infection (see below), a significant cytokine response, including marked activation of the antiviral ISG response, was observed in the heart at 4 dpi (Fig. 3F). Type I interferons IFNα and IFNβ, but not IFNγ, were increased, accompanied by increases in RIG-I (18-fold), OAS1, OAS2, CXCL10 (7-fold), CXCL11, CCL2, and IL-1β. There were no significant changes in IL-10, TGFβ or TNFα in the heart.

### Viral protein expression in lung but not heart

To test whether SARS-CoV-2 was directly infecting the heart, we probed for viral nucleocapsid expression in whole heart and lung homogenates by western blot or by immunofluorescence in lung or myocardial tissue slices(Fig. 3G,3H). Western blot showed no detectable viral nucleocapsid protein in myocardial lysates at 4 dpi, while two bands were readily detected in infected lungs (Fig 3G). Widespread nucleocapsid expression was evident in the lung at 4 dpi (Fig. 3H, left panel) but was absent in the myocardium (Fig. 3H, right panel). Consistent with protein expression, quantitative measurement with qPCR also revealed significant viral RNA expression in the infected lung at 4 dpi, while no viral RNA was detectable in heart or mock infected lung (Fig. 3I). These data suggest that the cardiac effects of SARS-CoV-2 infection are indirectly linked to an end organ or systemic inflammatory response rather than by direct viral infection and replication in cardiomyocytes.

### Effects of viral dsRNA activation of innate immune response on cardiac electrophysiology in guinea pigs

The strong activation of antiviral interferon signaling, in the absence of direct evidence of cardiac viral infection, begged the question of whether the ECG phenotype might be indirectly linked to the innate immune response to circulating viral RNA or myocardial viral infiltrates(*13*). Hence, we sought to determine if a viral dsRNA mimetic (Polyinosinic:polycytidylic acid; PIC) was sufficient to mimic the cardiac electrophysiological phenotype. We utilized naïve guinea pigs for a one-time direct myocardial injection of PIC into the left ventricular free wall. This species was used because of the close correspondence of its electrophysiological profile to humans, including a long action potential plateau, and repolarizing currents dominated by the rapid and slow components of the delayed rectifier potassium channels (*47–49*), thereby enabling a closer analysis of repolarization. Remarkably, bradycardia, sinus pauses, and AV nodal dysfunction were observed in PIC-treated guinea pigs, but not vehicle-injected controls (Fig. 4A-C). Mean RR was significantly increased by PIC treatment compared to controls (225.9±7.7ms in control vs 268.0±8.7ms in PIC, p<0.05) (Fig. 4B). The incidence of sinus pauses in PIC group (RR>RRmean+2SD) increased by 17-fold (15.9±3.7 in PIC vs 0.9±0.7 in control, p<0.05) (Fig. 4C). PIC treatment evoked a cardiac innate immune response, with elevated expression of OAS1, CCL2, TGFβ, IL-1β, TNFα, and Casp3 mRNAs (Fig.4D-I).

**Fig. 4.**
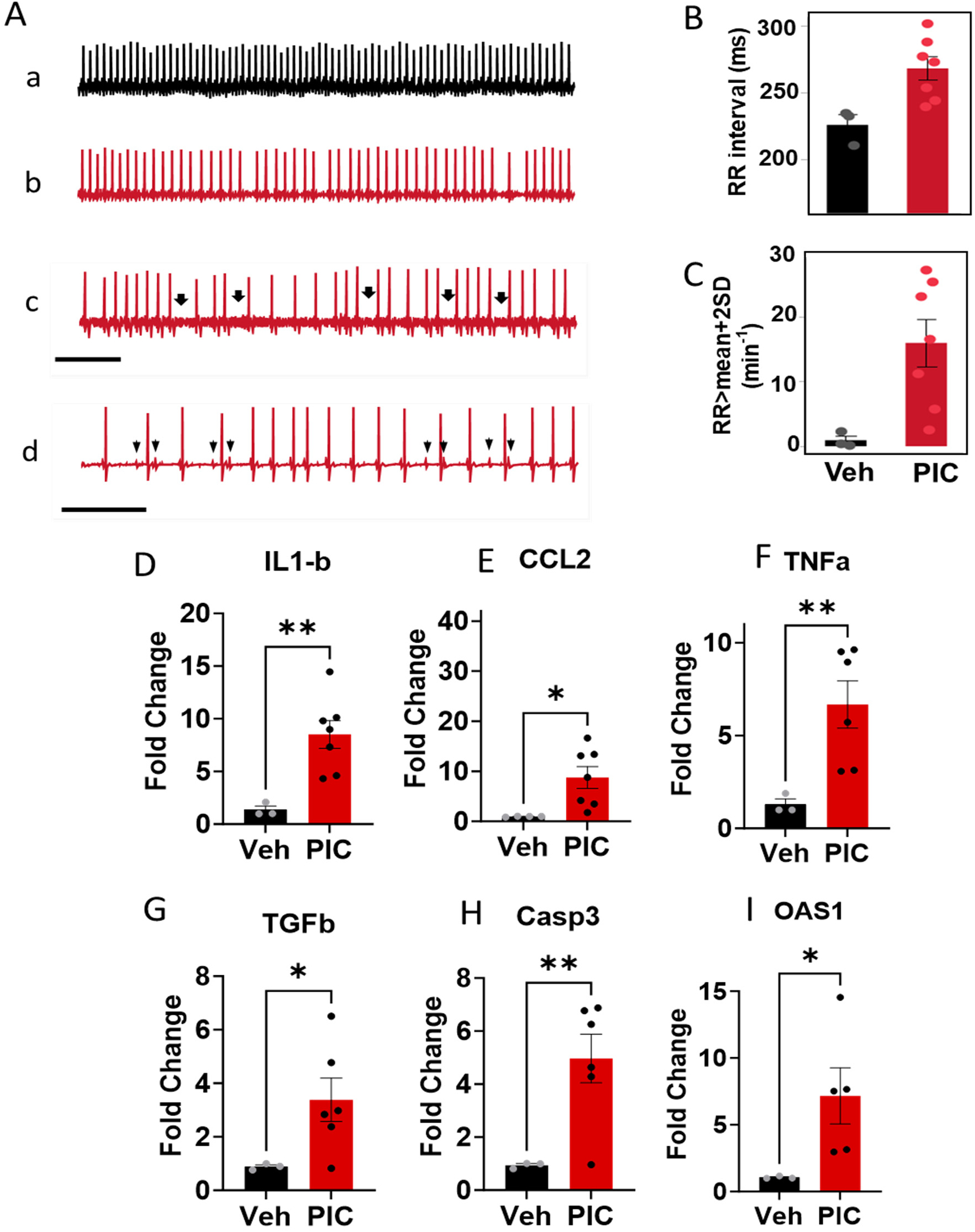
dsRNA induces cardiac arrhythmias and innate immune responses in the absence of viral infection in guinea pigs. **(A)** Representative ECG traces showing regular cardiac rhythm in a vehicle-injected control (a) and bradycardia (b), sinus pauses (c; marked by arrows), and AV block (d) after cardiac injection of polyinosinic-polycytidylic acid (PIC) in guinea pigs (arrowheads in d indicate abnormal P waves; horizontal bar equals 2 sec). **(B-C)** Summary data showing increase in RR Interval **(B)** and sinus pauses **(C)** in PIC-injected guinea pigs. **(D-I)** Increased expression of innate immune response genes induced by PIC, expressed as fold-change from vehicle-injected controls. Welch’s two tailed t-test was performed using alpha <0.05. * p<0.05, ** p<0.005

### Innate Immune Responses to dsRNA in human A549 cells and human induced pluripotent stem cell - derived cardiomyocytes

PIC treatment was used to examine the effects of activating the viral RNA-dependent innate immune response, independent of viral infection/replication, on interferon responses, cytokine release, and function. In a human lung epithelial cell line, A549, representing a frontline target of SARS-CoV-2, PIC treatment (added together with an empty adenovirus, AdV, to aid PIC uptake; see methods) activated the RNA pattern recognition pathway and the interferon response. We observe IRF3 and Stat1 phosphorylation, as well as increases in Stat1, IFNβ1 and OAS1,2 and 3 protein expression (Fig. 5A). In addition, RNA degradation was elevated by PIC, consistent with activation of RNase L (Fig. 5B). PIC also activated interferon signaling in hiPSC-CM monolayers, with increases in IRF9, MX1, OAS2 and 3 isoforms, and increases in STAT1 protein expression and phosphorylation (Fig. 5C).

**Fig. 5.**
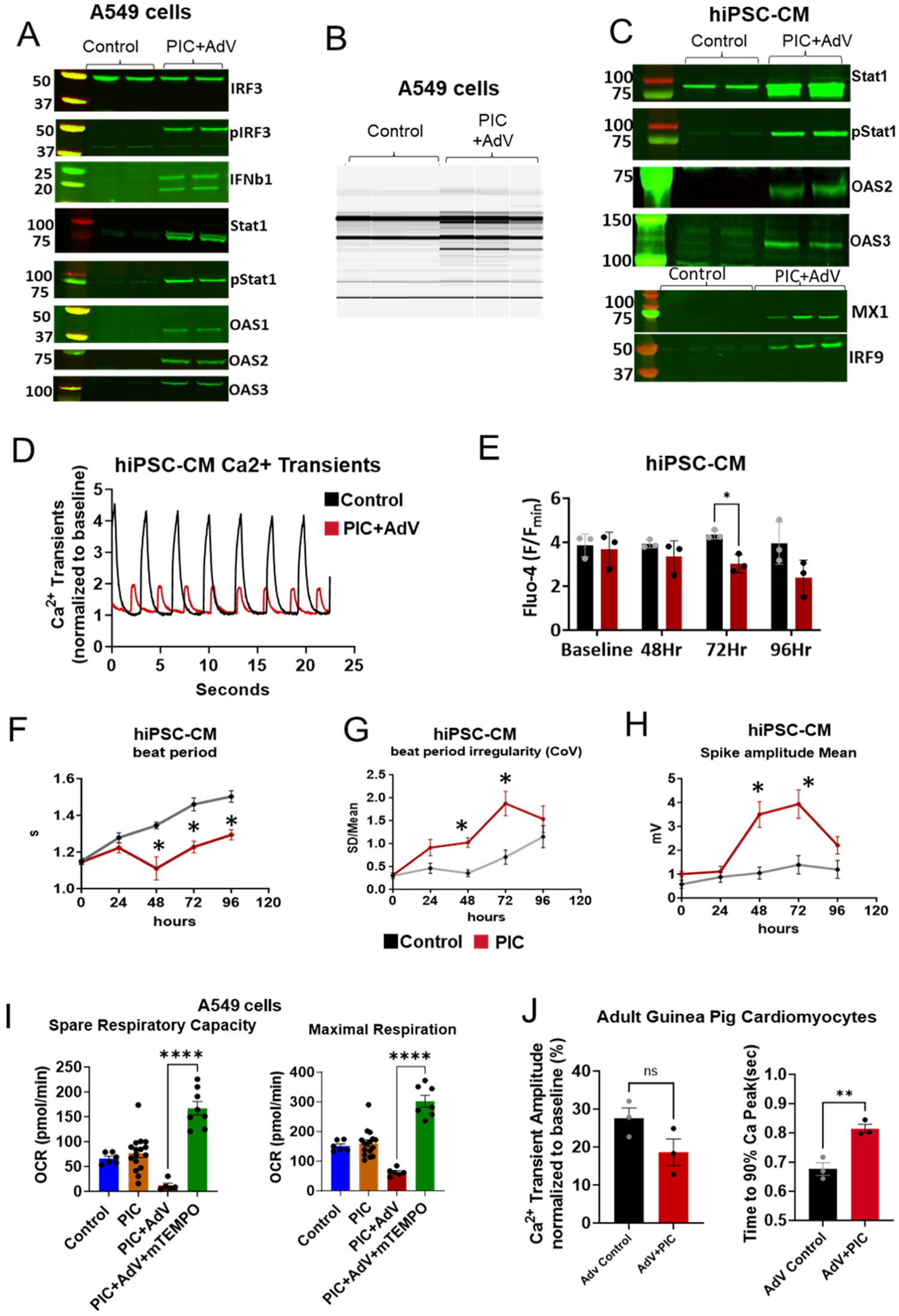
PIC increases expression of Type-I interferon signaling pathway proteins, activates RNA degradation, and alters electrophysiological and mitochondrial function. **(A-B)** Activation of Type I interferon signaling pathway proteins (A) and RNAse activation (B) in A549 lung epithelial cells. **(C)** Increase in interferon signaling pathway protein expression after PIC treatment in hiPSC-CMs. **(D)** Example Ca^2+^ transient traces show PIC (red trace) reduced Ca^2+^ transient amplitude in hiPSC-CMs compared to control (black trace). **(E-H)** PIC significantly reduced Ca^2+^ transient peak amplitude **(E)** by the 4^th^ day of treatment in monolayers of hiPSC-CMs (*p<0.03). Beat period was significantly lowered by days 2, 3 and 4 **(F)**(*p<0.001) and beat period irregularity was also significantly increased on day 3 after PIC treatment **(G)**(*p=0.0002). Mean spike amplitude was significantly increased by days 2 and 3 after PIC treatment **(H)** (p=0.02 and 0.03 respectively). Significance was estimated with a 2way ANOVA and Experiments were done thrice with different hiPSC-CM differentiation batches. **(I)** Mitochondrial Oxygen Consumption Rate (OCR; Seahorse XF96 assay) was suppressed after PIC+AdV treatment in A549 cells, including a decrease in both Maximal Uncoupled Respiration and Spare Respiratory Capacity. The mitochondrial deficiency was prevented by mitoTEMPO treatment, implicating mitochondrial ROS. Note: In A549 cells, PIC treatment alone elicited a weak interferon response that was markedly accentuated by co-addition of an empty replication-incompetent adenovirus vector, AdV (see Supplemental Fig. 1 and methods). AdV alone did not activate the immune response and was not cytotoxic, so subsequent immune challenges were performed with the combination of PIC+AdV in A549 cells. However, in hiPSC-CMs, PIC alone was enough to mount a substantial immune response with STAT1 and pSTAT1 increase (Supplemental Fig. 2) and addition of AdV did not significantly increase the immune response. Therefore, the experiments in hiPSC-CMs were generally performed with PIC alone, unless specifically labeled as PIC+AdV in the figures. **(J)** Ca^2+^ transient analysis from cardiomyocytes from guinea pig hearts injected with PIC+AdV or AdV alone. Unpaired t-test was used to determine significance with p<0.05. IRF3: interferon response factor 3, pIRF3: phospho-IRF3, IFNb1: interferon β1, Stat1: signal transducer and activator of transcription 1, pStat1: phospho-Stat1, OAS1(2,3): 2′-5′-oligoadenylate synthase 1(2,3), MX1: MX dynamin like GTPase 1, IRF9: interferon response factor 9.

To assess the effects of dsRNA innate immune activation on cardiac excitation-contraction coupling, we measured Ca^2+^ transients in hiPSC-CM monolayers treated with PIC+AdV. Innate immune activation resulted in a decrease in Ca^2+^ transient peak amplitude 48-96 hours after PIC+AdV treatment (Fig. 5D and 5E). PIC increased beating rate, beat period irregularity and spike amplitude compared to controls (Fig. 5F-H). PIC+AdV markedly suppressed both basal and maximal (uncoupled) oxygen consumption rate (OCR) in A549 cells, nearly eliminating spare respiratory capacity. (Fig 5I). The bioenergetic dysfunction after the immune challenge was mediated by excess mitochondrial ROS, as pretreatment with the mitochondrially-targeted antioxidant mitoTEMPO(*50, 51*) enhanced basal, maximal and spare respiratory capacity to levels even higher than the control group. Cardiomyocytes isolated from adult guinea pigs myocardially-injected with PIC+AdV show suppressed Ca^2+^ transients and significantly increased Ca^2+^ transient duration at 90% of decay (CaD90) (Fig. 5J).

In hiPSC-CM, Janus kinase/signal transduction and transcription activation (JAK/STAT) inhibition with 1µM ruxolitinib suppressed Type-I interferon protein expression (STAT1, pSTAT1, MX1, IRF9) and cytokine responses induced by PIC (Fig. 6A and B). We also analyzed the profile of secreted cytokines in media from hiPSC-CM subjected to the immune challenge using a multiplex antibody array (Proteome Profiler, R&D Systems). PIC induced the secretion of a wide variety of cytokines from the cardiomyocytes, including a >2-fold increase in 36 of the 105 cytokines assayed (Fig.6C and Supplementary Table 3). The most abundant cytokine secreted by the hiPSC-CM was CXCL10 (IP-10), which increased in the media ∼90-fold. This finding suggests that myocytes themselves could contribute to the whole heart increase in CXCL10 expression shown earlier (Fig. 3F). Interestingly, CXCL10 was recently linked to the cytokine storm in adults infected with SARS-CoV-2(*44, 46*) and is a biomarker of multisystem inflammatory syndrome and left ventricular dysfunction in children with COVID-19(*43*). Several other cytokines thought to contribute to the cytokine storm in patients with COVID-19 were secreted by hiPSC-CM after the PIC immune challenge, including IL-6 (10.4-fold +/-7.2 SEM)(*45*), VEGF (7.2-fold +/-2.6 SEM)(*52*), HGF (6-fold +/-2.9 SEM)(*53*), CXCL5 (17.3 fold +/-5.3 SEM)(*54*), IL-8 (11.8-fold +/- 4.2SEM)(*55*), and CXCL1 (6.3 fold+/- 1.8 SEM)(*55*), potentially representing local mediators of myocardial inflammation, along with CCL5 (17.8-fold +/- 5.5 SEM), which was associated with COVID-19 severity(*56*). 1µM ruxolitinib suppressed cytokine secretion and brought it back close to control levels. To understand the extent of oxidative stress in the cellular innate immune response, we also treated hiPSC-CM with a mitochondrially-targeted antioxidant, mitoTEMPO (1µM). MitoTEMPO did not suppress Type-I interferon protein induction, nor the cytokine response induced by PIC (Fig. 6A). We next assessed mitoTEMPO and ruxolitinib treatment on Engineered Heart Tissues (EHTs). 1 µM ruxolitinib lowered Type-I IFN responses, with significant decreases in STAT1, pSTAT1, MX1 and IRF9, while 1 µM mitoTEMPO treatment had no statistically significant effect, although the IFN response tended to be lower (6D and E).

**Fig. 6.**
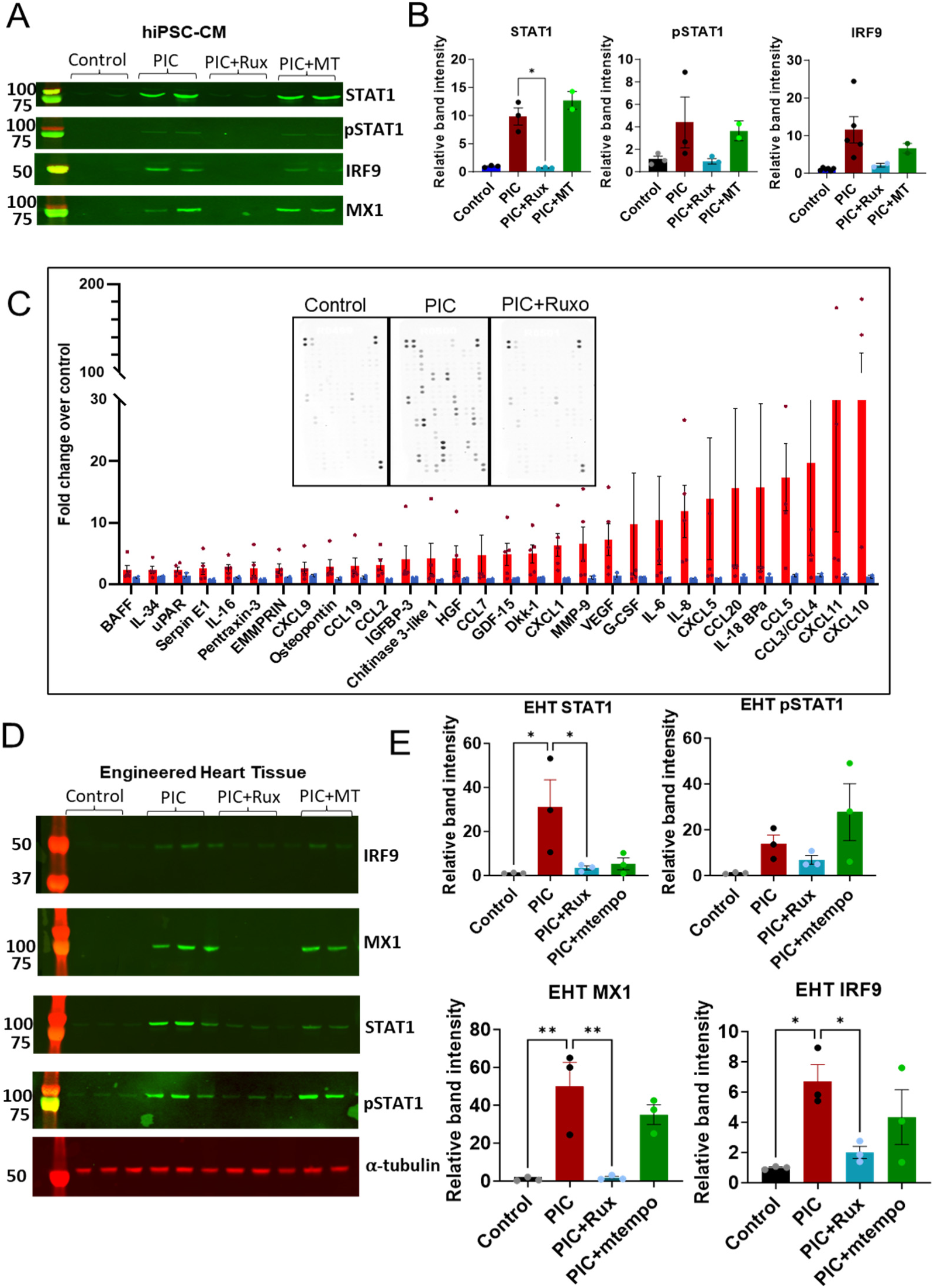
JAK/STAT inhibition, but not mitochondrial antioxidant treatment, suppresses interferon responses in hiPSC-CMs. **A-B)** Inhibition of JAK/STAT signaling with Ruxolitinib (1μM) suppressed PIC-mediated induction of the interferon response signaling proteins STAT1, pSTAT1, IRF9 and MX1 in hiPSC-CM monolayers. Bar graphs show relative band intensity normalized to Ponceau-stained total protein loading. **C)** Cytokine responses induced by PIC were greatly suppressed by Ruxolitinib (Rux) treatment in hiPSC-CMs. Inset is an example array showing Control, PIC and PIC+Ruxolitinib. **D)** Inhibition of JAK/STAT signaling in hiPSC-derived Engineered Heart Tissues (EHTs) with Ruxolitinib, but not mitoTEMPO (MT), significantly decreased the interferon response signaling proteins STAT1, pSTAT1, IRF9 and MX1. **E)** Relative band intensity normalized to Ponceau total protein loading for EHTs (2-way Anova, with p<0.05. Lysates loaded for the western blot in D were from 3 different EHTs).

### Effects of JAK/STAT inhibition or mitochondrial ROS scavenging on COVID-19 induced pulmonary and cardiac dysfunction

The effects of activation of dsRNA-triggered innate immune signaling on hiPSC-CM function provided motivation to test the therapeutic potential of systemic inhibition of JAK/STAT signaling in the COVID-19 hamster model. Ruxolitinib is one of several clinically utilized JAK/STAT inhibitors (specifically, a JAK1,JAK2 inhibitor) that are potent anti-inflammatory and immunosuppressive agents commonly prescribed for autoimmune disease (*57, 58*). A clinical trial for its use in COVID-19 was initiated, but prematurely truncated, by the sponsor due to low statistical power (*59*) and FDA approval of the alternative JAK/STAT inhibitor baricitinib for combination therapy, which reduced mortality in hospitalized COVID-19 patients(*60*). Oxidative stress also contributes to the innate immune response and the cytokine storm, so antioxidant interventions have been proposed as potential treatments for COVID-19 complications (*61–63*). To assess the impact of inhibiting JAK/STAT or mitochondrial ROS on pulmonary and cardiac functional parameters in the hamster COVID-19 model, we implanted osmotic pumps for chronic intraperitoneal delivery of ruxolitinib (2 mg/kg/day) or mitoTEMPO (1.1 mg/kg/day) 4 days prior to SARS-CoV-2 infection and continued until 10 dpi, covering the acute phase of infection. Pulmonary function and ECG analyses were carried out as described above. The results for the 2 treatments are shown (Fig. 7) superimposed on the data for the untreated SARS-CoV-2 infected group (replotted from Figs. 1 and 2).

**Fig. 7.**
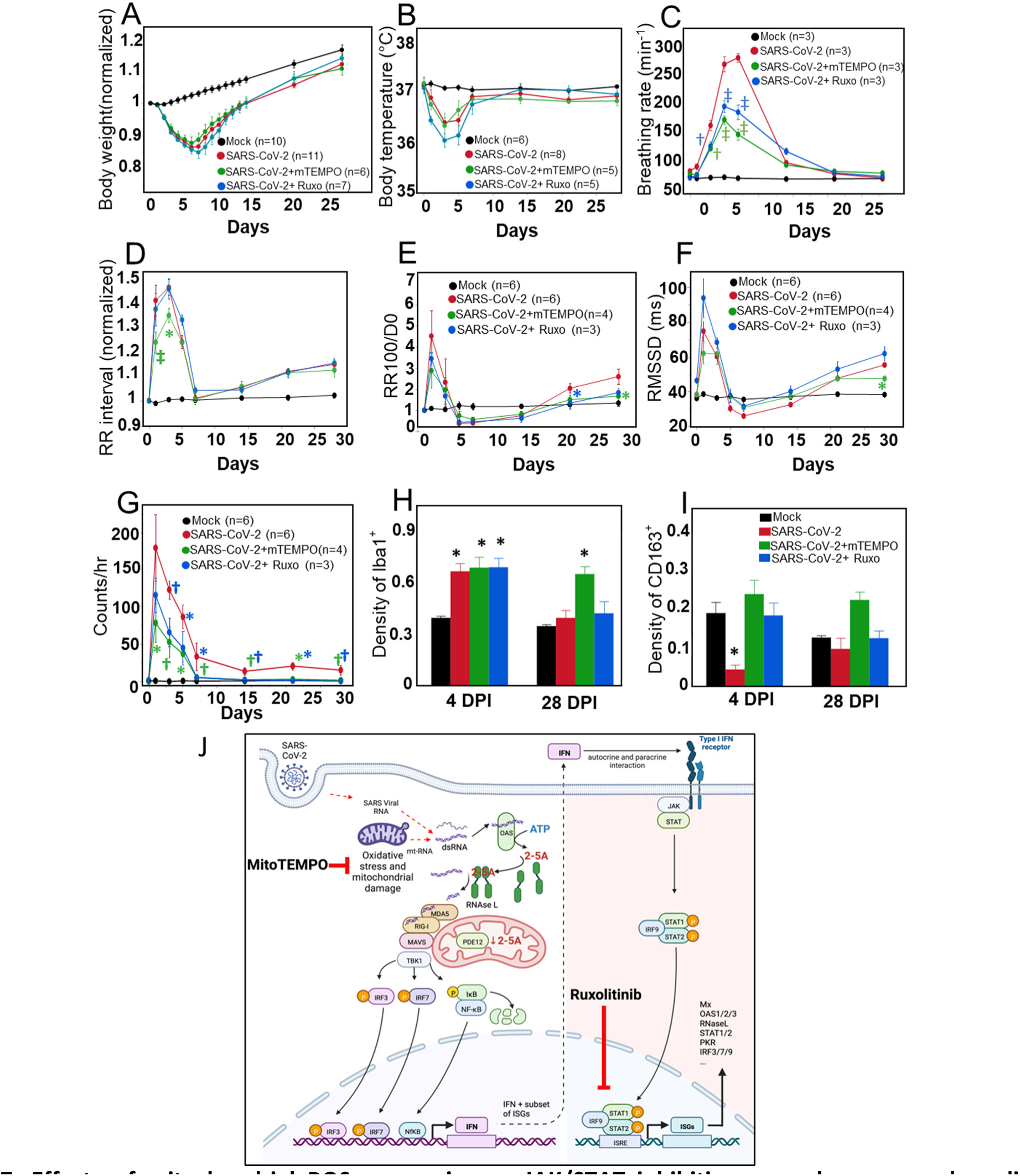
Effects of mitochondrial ROS scavenging or JAK/STAT inhibition on pulmonary and cardiac sequelae of COVID-19. **(A-B)** COVID-19-induced changes in body weight **(A)** or temperature **(B)** were not significantly altered by mitoTEMPO (mTEMPO) or Ruxolitinib (Ruxo) treatments. **(C)** SARS-CoV-2-induced tachypnea was significantly attenuated by both mTEMPO and Ruxo treatments. † p<0.005, ‡ p<0.0001. **(D)** mTEMPO, but not Ruxo, attenuated the increase in RR interval at 1 and 3 dpi. After returning to baseline level at 7 dpi, both treatments showed no impact on the redeveloped bradycardia. * p<0.05, ‡ p<0.0001 **(E-F)** The early spike in sinus pauses **(E)** and RMSSD **(F)** at 1 dpi were not significantly decreased by either treatment. Both Ruxo and mTEMPO attenuated the late effects of SARS-CoV2 infection on sinus pauses at 21 or 28 dpi, respectively (*p<0.05). Only mTEMPO significantly suppressed RMSSD at 28 dpi (*p<0.05). **(G)** Both treatments significantly attenuated the incidence of AC block from 1 – 7 dpi and abrogated the sustained increase in AV block occurrence at 14-28 dpi. (*p<0.05, † p<0.01, ‡ p<0.0001; compared to SARS-CoV-2 alone, blue symbols are Ruxo treatment, green symbols are mTEMPO treatment; see Supplementary Table 4 for additional within group comparisons with Day 0) **(H)** IBA1^+^ macrophage density in the cardiac conduction system region increased in all SARS-CoV-2-infected groups regardless of treatment at 4 dpi and returned to baseline level at 28 dpi, except in the mTEMPO treated group (*p<0.05). **(I)** Treatment with either mTEMPO or Ruxo prevented the decrease in CD163+ macrophage density in the conduction region at 4 dpi. *p<0.05 compared to Mock. 3-11 hamsters per group per parameter were analyzed. n=3 for groups of 4 dpi and n=4 for groups of 28 dpi. Each heart has 3 repeats. **(J)** Schematic of molecular mechanisms of COVID-19 and dsRNA induced IFN response. Ruxolitinib suppresses IFN stimulated response and mitoTEMPO suppresses mitochondrial damage and oxidative stress. (Created with BioRender.com).

**Fig. 8.**
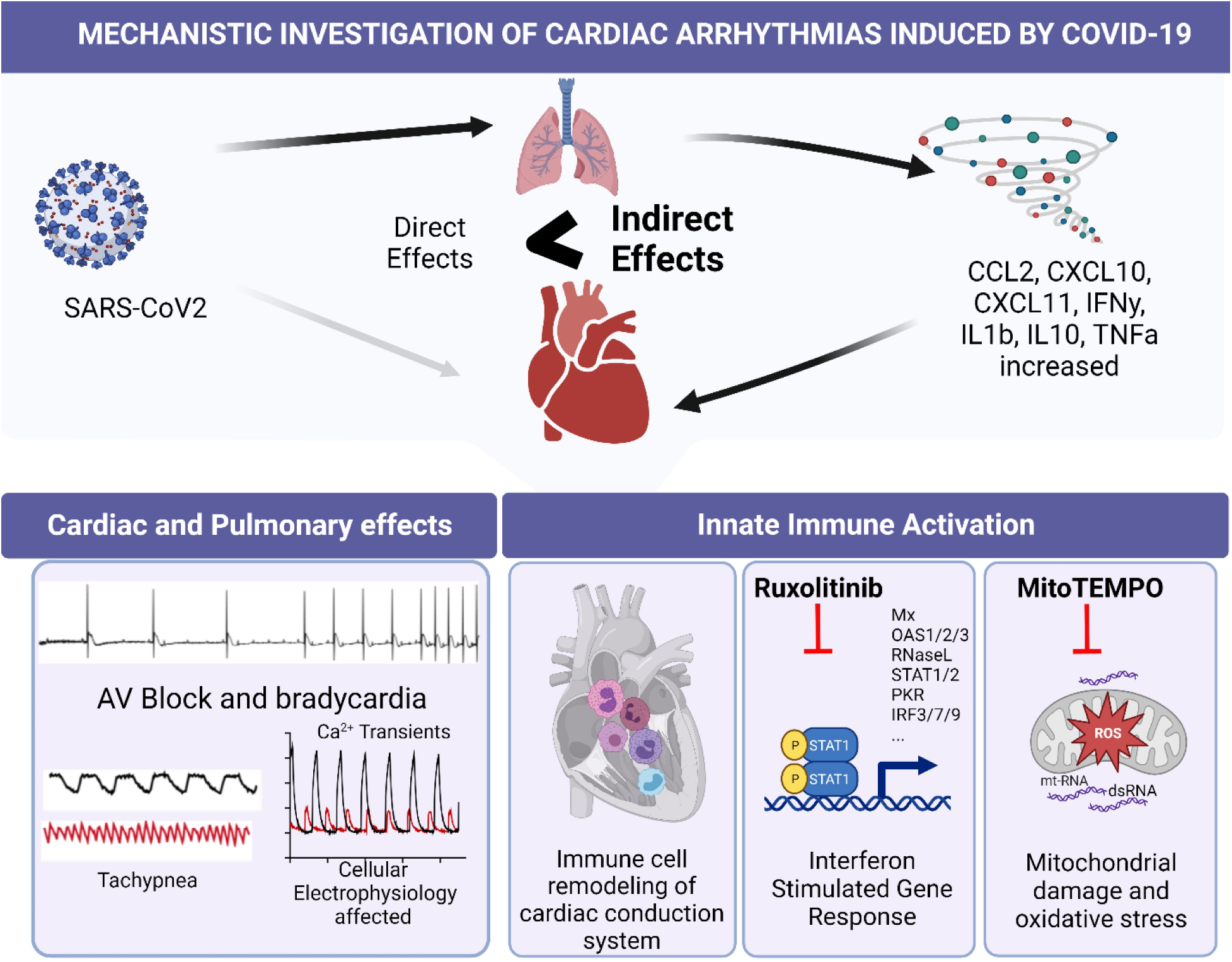
Graphical Abstract. We performed a mechanistic investigation of cardiac arrhythmias induced by SARS-CoV-2. We demonstrate that persistent cardiac conduction system dysfunction associated with COVID-19 arises indirectly from oxidative stress and excessive activation of cardiac innate immune responses during infection (Created with BioRender.com).

Ruxolitinib or MitoTEMPO treatments had minimal effect on the decreases in body weight or body temperature observed after SARS-CoV-2 infection (Fig. 7A and B); however, both treatments significantly inhibited the tachypnea associated with COVID-19. Ruxolitinib treatment decreased the peak breathing rate by 50% and 40% at 5 and 7 dpi, respectively, compared to SARS-CoV-2 infection alone. MitoTEMPO treatment decreased the peak breathing rate by 62% and 108% at 5 and 7 dpi, compared to untreated infected animals. MitoTEMPO treatment was more effective than ruxolitinib treatment at suppressing tachypnea, and the recovery to baseline was faster (Fig.7C). Interestingly, mitoTEMPO treatment improved the cardiac electrophysiological phenotype, blunting the increase in RR interval over the first 7 dpi (Fig. 7D). Although mitoTEMPO was only administered during the acute phase, its effect extended to post-infection phase. The prolonged response of RR100 and RMSSD to SARS-CoV-2 infection was suppressed, and no significant difference in RR100 or RMSSD at 21 and 28 dpi was observed, compared to 0 dpi (Fig. 7E and F). Ruxolitinib had no obvious effects on sinus bradycardia and autonomic dysfunction associated with COVID-19; however, both mitoTEMPO and ruxolitinib prevented AV block in the post-infection phase from 7 – 28 dpi, where the incidence of AV block remained higher than baseline in the group with SARS-CoV-2 infection alone (Fig. 7G). We next examined macrophage densities and found that the increase in Iba1+ macrophages in the CCS in the COVID-19 group at 4 dpi was not prevented by either of the treatments (Fig.7H); however, both treatments prevented the early decline in CD163+ macrophages (resident or M2-type macrophages) (Fig.7I). At 28 dpi, Iba1+ and CD613+ macrophage counts reverted to the levels of uninfected controls in the all of the SARS-CoV-2 groups except the mitoTEMPO treated.

## Discussion

This study provides a comprehensive time course and mechanistic investigation of the short- and long-term cardiac arrhythmias induced by COVID-19 in an animal model susceptible to infection with human SARS-CoV-2. The major findings are that 1) COVID-19 causes substantial CCS dysfunction, including acute bradycardia, sinus node dysfunction and AV nodal block that follows a triphasic time course, consisting of a transient increase in arrythmias during the peak of lung infection, partial reversion to normal, and a late redevelopment of arrhythmias; 2) dsRNA-triggered activation of the innate immune response, in the absence of viral infection, mimics the electrophysiological phenotype in vivo and alters the cytokine secretome and excitation-contraction coupling functions of human cardiomyocytes; 3) remodeling of macrophage phenotypes of the CCS and ventricles occurs during COVID-19; and 4) inhibition of interferon signaling or suppression of mitochondrial oxidative stress blunts both the pulmonary and cardiac electrophysiological effects of COVID-19 (see schematic, Fig. 7J).

### Cardiovascular consequences of COVID-19

During the COVID-19 pandemic, several studies reported that the risk of new onset cardiometabolic disease increased dramatically after SARS-CoV-2 infection. For example, a case-control study of >400k patient medical records in the UK health system showed large increases in relative risk of diabetes, thrombosis, heart failure, and cardiac arrhythmias that spiked early, but persisted for months after the index infection(*1*). Similarly, a worldwide survey reported that arrhythmias occurred in ∼18% of COVID-19 patients(*64*), with the majority developing atrial arrhythmias (fibrillation or flutter), ∼23% developing bradyarrhythmia (sinus pauses or AV block), and ∼20% showing ventricular tachyarrhythmias (VT or VF). A high incidence of arrhythmias was also reported in COVID-19 patients admitted to the intensive care unit, varying between ∼17%(*4, 65*) and 44% (*66, 67*), with many patients displaying AV block, bradycardia and relative bradycardia(*68–71*), associated with worse prognosis(*70*). In one study of COVID-19 patients with bradycardias, complete heart block was highly prevalent and pacemaker implantation was required in 76% of these cases(*72*). Arrhythmias involving impaired CCS activity are also significantly increased in patients suffering from long COVID syndrome (also known as post-acute sequelae of COVID-19 or PASC). These include chronotropic incompetence, increased heart rate variability or postural orthostatic tachycardia syndrome (POTS), which points to long-term dysautonomia or impaired function of the CCS(*73*). Since then, widespread vaccination has become available, but it is unclear how cardiovascular complications have been impacted at the population level. Nevertheless, it remains critically important to elucidate the mechanisms underlying acute and persistent virus-induced cardiac arrhythmias, given the ongoing evolution of SARS-CoV-2, the inability of the vaccines to prevent new infections, and the ongoing potential of a variety of respiratory viruses to induce CCS dysfunction(*74*).

In the context of these human studies, our results show that pulmonary SARS-CoV-2 infection in hamsters is a robust model to study the bradyarrhythmias and CCS dysfunction associated with COVID-19, in particular, severe types of AV block. Spontaneous atrial tachyarrhythmias, which were prominent in humans with COVID-19 (*75*), were not generally observed in this model, but there are two factors that might account for this. First, spontaneous reentrant arrhythmias, such as atrial fibrillation or flutter, are rarely observed in small animals owing to the small mass (and conduction path length) of the heart. Thus, most studies utilize programmed stimulation to examine atrial arrhythmia inducibility(*76*). Second, age is a major cofactor associated with atrial fibrillation in the general population, and in COVID-19 patients (*75*). Our young adult hamsters lack increased fibrosis and other co-morbidities of age that predispose to atrial fibrillation. Although the hamster model faithfully reproduces a subset of COVID-19-associated arrhythmias, the inflammatory remodeling of the CCS and the ECG changes that we observed are indicators of altered excitability and conduction that could be a substrate for more complex atrial or ventricular tachyarrhythmias under stress or in the presence of co-morbidities. Moreover, our data show that the hamster model displays long-term effects on sympathovagal balance, which is disrupted in long COVID syndrome(*77*).

### Cardiac innate immune responses underlie cardiac arrhythmias

The mechanisms underlying the increased arrhythmia incidence in COVID-19 are unknown. Early studies of autopsy samples suggested that replicative SARS-CoV-2 RNA is present in the hearts of COVID-19 patients (*11, 12*). This view has given way to the consensus that viral RNA may be present (*13*), but little direct myocardial infection occurs(*21–24*). Nevertheless, evidence of cardiac inflammation is widely reported(*15–19*), supporting the systemic cytokine storm as a possible mechanism(*19, 78–80*). Consistent with many human studies, we found no evidence of viral protein expression in hearts of hamsters after nasal infection with SARS-CoV-2, which differs from a previous study reporting ∼10-15% of myocytes with positive antibody signals for viral spike or N protein (*81*). The reason for this discrepancy is unclear but could be related to SARS-CoV-2 strain or titer differences. Regardless, we show here that the cardiac arrhythmias, cytokine responses, and macrophage infiltration occur in the absence of cardiac viral protein expression, and can be mimicked by direct activation of the innate immune response with dsRNA. The opposing changes in the numbers of Iba1+ and CD613+ macrophages in the AV nodal region early after infection, paralleling CCS dysfunction, are particularly interesting. Tissue resident macrophages highly express CD163(*42*) and can influence cardiac conduction by direct electrical coupling with nodal myocytes(*40*). They also broadly participate in myocyte mitochondrial quality control by removing damaged mitochondria through an exopher mechanism(*38*). In contrast, monocyte-recruited inflammatory type macrophages can elicit atrial fibrillation(*82*). We propose that the observed CCS remodeling and altered local crosstalk between myocytes and immune cells is responsible for impaired function. It should be noted that we detected a plethora of chemoattractant cytokines and growth factors released from human cardiomyocytes after activation of the innate immune response. These could act in an autocrine or paracrine manner to alter myocyte function, or recruit additional immune cells, in turn, releasing a different complement of factors, underlining the complexity of the extracellular milieu.

### Cytokine Profiles

Increased cytokines and interferon-stimulated gene expression were present in both lung and heart tissues of SARS-CoV-2-infected hamsters (Fig. 3). Several of these have been associated with severity of disease in human COVID-19 patients, as well as in other arrhythmogenic diseases. For example, increases in CCL2 and the receptor CCR2 were observed in a murine arrhythmogenic model, along with other inflammatory markers(*83*), and the CXCL12-CXCR4 axis was identified as a key mediator of atrial fibrillation(*84*). In the latter case, CXCL12 was upregulated and treatment with a CXCR4 (receptor for CXCL12) antagonist reduced inflammatory cells and markers in the atrial region and suppressed atrial fibrillation. CXCL10 and CXCL11 have been implicated as mediators of the cytokine storm in SARS-CoV-2 infection(*43, 44, 46*), but they are also recognized as inflammatory mediators in ischemic heart disease and heart failure(*85, 86*). Elevated IL-6 was associated with bradycardia (*87*), conduction abnormalities, and atrial fibrillation (*88, 89*) in COVID-19 patients.

### dsRNA activation of innate immune responses in the absence of virus

Interestingly, direct injection of guinea pig hearts with the dsRNA mimetic PIC resulted in arrhythmias that were similar to those observed in the COVID-19 hamsters, i.e., bradycardia, long sinus pauses and AV nodal block. PIC also increased expression of innate immune response genes, supporting the hypothesis that Type-I IFN stimulation in the absence of viral replication is sufficient to phenocopy the in vivo electrophysiological phenotype of COVID-19. To better understand which secreted factors might underlie the cellular effects of innate immune activation, we exposed human iPSC-derived cardiac myocytes or engineered heart tissues to PIC and observed pronounced Type-I interferon signaling responses. Even highly purified cardiomyocyte monolayers (98% enriched myocytes) released an abundance of cytokines and growth factors, indicating that myocytes themselves can contribute to the antiviral response, as many of these corresponded to those observed in COVID-19. For example, CXCL10 and CXCL11, which are upregulated in COVID-19 patients (*44, 46*), were markedly increased in the medium after PIC activation in hiPSC-CMs, together with more than 20 other cytokines. PIC also impacted adult guinea pig and hiPSC cardiomyocyte function, blunting cytosolic Ca^2+^ transients and affecting decay kinetics. Beat irregularity and frequency changes were also observed in hiPSC-CM monolayers. Similarly, depressed Ca^2+^ transient amplitude and increased spontaneous Ca^2+^ release events were previously reported in hiPSC-CMs and rat ventricular myocytes exposed to serum from COVID-19 infected patients (*90*), possibly implicating cytokines as mediators of rhythm disturbances that could translate to higher organ level cardiac arrhythmias. Further studies are needed to discern which cytokine-receptor combinations might be critical to the CCS remodeling underlying the electrophysiological phenotype in COVID-19.

### Therapeutic implications

The expression of RIG-I, OAS1 and OAS2 genes were also significantly increased in the lungs and hearts of SARS-CoV-2-infected in hamsters. The 2’,5’-oligoadenylate (2-5A) synthase (OAS) family of proteins are key upstream pattern recognition receptors in the antiviral interferon response. Notably, a gene variant in the OAS1/2/3 gene locus is associated with SARS-CoV-2 susceptibility and alters the antiviral response (*91–93*). Downstream of OAS, the dsRNA sensors RIG-I and MDA5, together with the mitochondrial antiviral signaling protein (MAVS), are localized in a complex on the outer mitochondrial membrane (*94*) that transduces the signal to activate early interferon response genes. Interestingly, OAS is overexpressed in post mortem hearts of patients who died of COVID-19, along with alterations of mitochondrial genes, even though no traces of viral gene expression were present, indicative of a strong cardiac innate immune response and perturbation of mitochondrial energetics (*22*). Mitochondrial impairment and oxidative stress are recognized intermediaries of the inflammatory response to viral infections (*95*) and increased levels of circulating mtDNA are correlated with severity and mortality in COVID-19 (*96*). Mitochondrial dysfunction itself could alter the antiviral interferon response by modifying 2-5A levels (*97*) or by triggering the release of mitochondrial DNA into the cytoplasm to activate the cGAS-STING pathway (*95, 98*), which was shown to contribute to endothelial cell damage during SARS-CoV-2 infection (*99, 100*).

Given the distinct inflammatory cascade signature promoted by SARS-CoV-2 and the induction of interferon-stimulated gene expression, we targeted two potential mechanisms of action: inhibition of JAK/STAT signaling (with ruxolitinib) and mitochondrial reactive oxygen species (with mitoTEMPO). Ruxolitinib is FDA approved to treat certain conditions, primarily myelofibrosis, polycythemia vera, and acute graft-versus-host disease (GVHD). A clinical trial for therapeutic potential of ruxolitinib for COVID-19 yielded promising results of reduced mortality and improved secondary outcomes, but the study was terminated due to lack of statistical power (*59*). Other JAK inhibitors have also been used, such as baricitinib, which showed promising preliminary results (*101*). MitoTEMPO has been used in preclinical models to mitigate ROS-induced adverse cardiac effects of heart failure, diabetes or aging (*50, 102, 103*). Treatment with either ruxolitinib or mitoTEMPO significantly blunted the tachypnea associated with SARS-CoV-2 infection, with the latter being somewhat more effective. MitoTEMPO, but not ruxolitinib, also suppressed the early transient bradycardia after infection, as well as the long sinus pauses at 28 dpi. Interestingly, either ruxolitinib or mitoTEMPO eliminated the persistent AV block events present at 15-28 dpi in COVID-19 hamsters. Limited information is available about the efficacy of antioxidant therapies in COVID-19, but in vitro work showed that mitoquinol (MitoQ) or N-Acetyl Cysteine, which enhance antioxidant defenses, inhibited viral replication in monocytes infected with SARS-CoV-2 (*104*). In two small clinical studies, treatment with N-Acetyl Cysteine reduced the clinical complications and mortality of COVID-19 patients (*31, 105*).

### Perspectives

The present findings are the first to describe the time course and mechanisms of cardiac arrhythmias associated with COVID-19 in an animal model that reproduces a subset of those reported in human studies. The observed bradyarrhythmias and nodal dysfunction point to both acute and chronic remodeling of the CCS, caused by activation of interferon signaling and mitochondrial dysfunction. Intervening in these pathways prevented the adverse cardiac and pulmonary effects, but additional investigation will be needed to determine the extent of permanent CCS remodeling after SARS-CoV-2 infection and if this provides the substrate for known electrophysiological abnormalities associated with long COVID syndrome.

## List of Supplementary Materials

1. Supplementary Figures-Supplementary Figure1-3
2. S1-Supplement Table 1-List of Antibody sources
3. S2-Supplement Table 2-List of qPCR Primers
4. S3-Supplement Table 3-Numerical data from Protein Array experiments
5. S4-Supplement Data File 4-Results of statistical analysis of comparisons between different treatment groups of hamsters.

