## Supplemental Data for "Innate Immune Activation and Mitochondrial ROS Invoke Persistent Cardiac Conduction System Dysfunction after COVID-19"

### Supplementary Materials

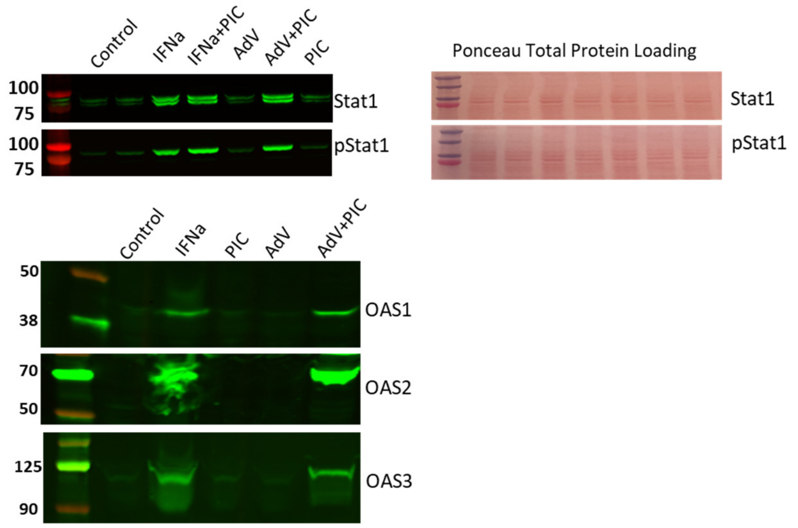

Supplementary Figure 1: In A549 cells, Poly(I:C) alone elicited a weak IFN response.

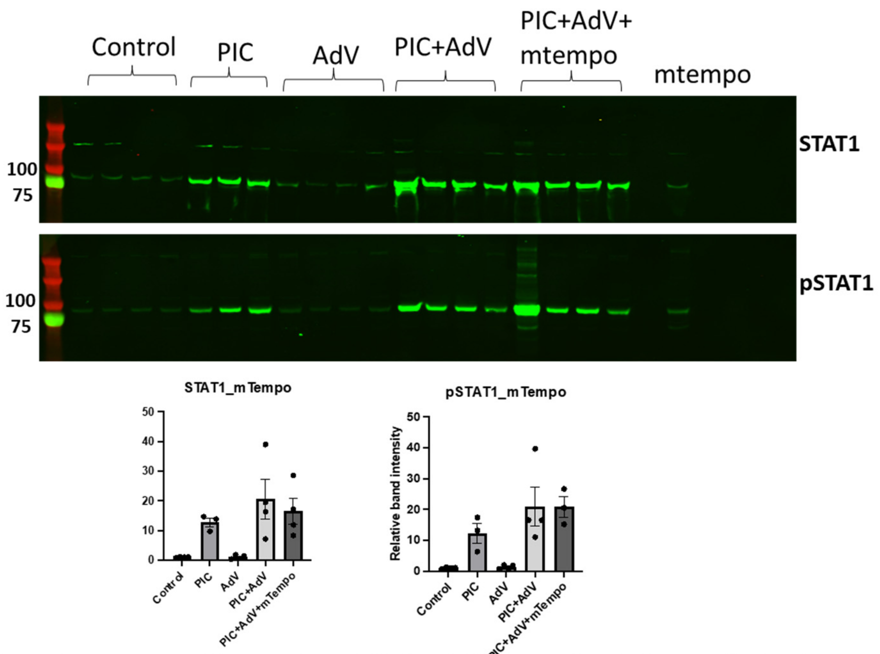

Supplementary Figure 2: In hiPSC-CMs, Poly(I:C) alone elicited a strong IFN response.

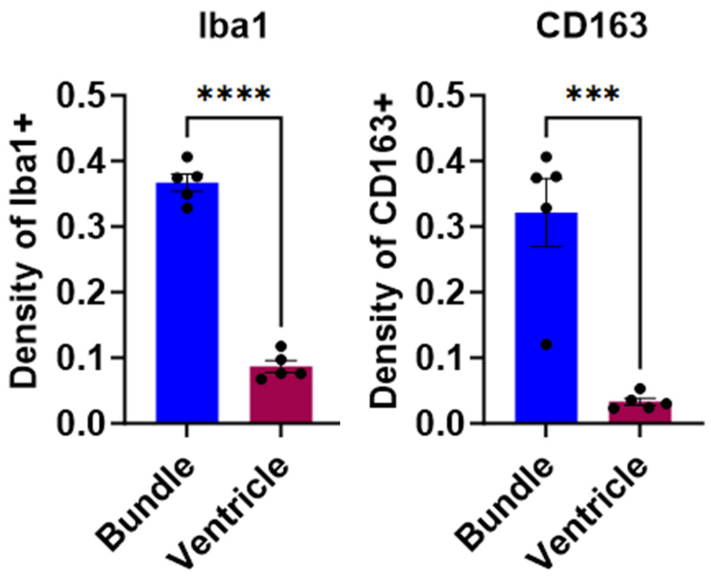

**Supplementary Figure 3:** Relative densities of Iba-1+ and CD163+ macrophages were higher in the AV/bundle region compared to ventricles (Iba-1+ macrophages: 0.37±0.01 in bundle vs 0.09±0.01 in ventricles, p<0.0001; CD163+ macrophages: 0.16±0.02 in bundle vs 0.03±0.01 in ventricles, p<0.005 in mock-infected control hearts.

Supplement Table S1- List of Antibody Sources

| Antibody | usage | host | Catalog # | Vemdor | dilution |
|---|---|---|---|---|---|
| IBA1 | IF | rabbit | A1527 | Abclone | 1:200 |
| Contactin2 | IF | Goat | AF-4439 | R&D | 1;50 |
| CD163 | IF | Rabbit | AB182422 | Abcam | 1:100 |
| SARS-CoV2 nucleoCapsid | IF/W | rabbit | 40143-R001 | Sino Biol. | 1:500/1:2000 |
| Anti-sheep, FITC | IF | Donkey | STAR88D488 | BioRad | 1:400 |
| anti-rabbit, TRITC | IF | Goat | A16123 | ThermoFisher | 1:800 |
| anti-Rabbit, 800CW | W | Goat | 926-32211 | Li-Cor | 1:20000 |
| Phospho-IRF-7 | W | rabbit | 12390S | Cell Signaling Technology | 1:1000 |
| IRF-7 | W | rabbit | 13014S | Cell Signaling Technology | 1:1000 |
| Phospho-STAT1 | W | rabbit | 8826 | Cell Signaling Technology | 1:1000 |

| | | | | | |
|---|---|---|---|---|---|
| STAT1 | W | rabbit | 14994 | Cell Signaling Technology | 1:1000 |
| IRF-9 | W | rabbit | 76684 | Cell Signaling Technology | 1:1000 |
| MX1 | W | rabbit | 37849 | Cell Signaling Technology | 1:1000 |
| OAS1 | W | rabbit | 14498 | Cell Signaling Technology | 1:1000 |
| OAS2 | W | rabbit | 54155S | Cell Signaling Technology | 1:1000 |
| OAS3 | W | rabbit | 41440S | Cell Signaling Technology | 1:1000 |

Supplement table S2- Hamster primers

| gene | Forward Primer | reverse primer |
|---|---|---|
| b-actin | CCA GAG CAA GAG AGG TAT T | TCG TTG TAG AAG GTG TGG |
| CCL2 | GAA AGA TCC CAG AGA AGA G | CTT GAG CTT GGT GAT GA |
| CXCL10 | GTG ACC TGT GGA TTG TTG | TTC TGG CTC TTC CTG TAT AA |
| CXCL11 | CGC CTC ATA CGG GAA ATG TAT | CAT CAG ACA CTC CCT GGT TTC |
| IFNa | TCC CAC CAA CTC ACT ATA C | CAA GAG GAT TCC GTG ATA TTT |
| IFNb | TAT CCC TGT CCA TCA ACT AC | CAC CTC CAT AGG CAT CTT |
| IFNy | TGT TGC TCT GCC TCA CTC AGG | AAG ACG AGG TCC CCT CCA TTC |
| IL-1b | CTG AAA GCT CTC CAT CTC | GCC ACA GGT ATC TTG TT |
| IL-6 | GGA CAA TGA CTA TGT GTT GTT AGA A | AGG CAA ATT TCC CAA TTG TAT CCA G |
| IL-10 | GGT TGC CAA ACC TTA TCA GAA ATG | TTC ACC TGT TCC ACA GCC TTG |
| OAS1 | TCT CCA AGG TGA TGA AGG | GCT GGT GAG ATT GTT AAG G |
| OAS2 | CAC CAT GAG AAG TAC AAT AAG | ATC AAA GGC TGG AAG TAG |
| OAS3 | GCT GCC CTC TAG TTA TG | GTC CTT GTT CTG TTG GA |
| RIG1 | GTG ACC TGT GGA TTG TTG | TTC TGG CTC TTC CTG TAT AA |
| TGFb | GGC TAC CAC GCC AAC TTC TG | GAG GGC AAG GAC CTT ACT GTA CTG |
| TNFa | TGA GCC ATC GTG CCA ATG | AGC CCG TCT GCT GGT ATC AC |
| Sars-CoV2 delta Spike | CCA CAA AAA CAA CAA AAG TTG G | TGA GAG ACA TAT TCA AAA GTG CAA |

Supplement Table S3- hiPSC-CM-Cytokine Array

| | Poly(I:C)(n=5) | SEM | +Ruxo(n=2) | SEM |
|---|---|---|---|---|
| Adiponectin/Acrp30 | 1.2085628 | 0.067297 | 1.084691 | 0.043779 |
| Angiogenin | 2.1054212 | 0.966669 | 0.91118 | 0.094981 |
| Angiopoietin-1 | 2.2785056 | 0.697605 | 0.925771 | 0.064425 |
| Angiopoietin-2 | 1.332314 | 0.106618 | 1.05127 | 0.140703 |
| Apolipoprotein A-1 | 1.4524768 | 0.074532 | 0.875808 | 0.103923 |
| BAFF/BLyS/TNFSF13B | 2.3284252 | 0.743817 | 1.080638 | 0.137438 |
| BDNF | 1.5601756 | 0.16595 | 1.047205 | 0.132103 |
| C-Reactive Protein/CRP | 1.3488904 | 0.082289 | 0.912717 | 0.070433 |
| CCL17/TARC | 1.4424288 | 0.061948 | 0.965699 | 0.222105 |
| CCL19/MIP-3 beta | 3.0288266 | 1.251598 | 1.113481 | 0.259478 |
| CCL2/MCP-1 | 3.1723178 | 0.915611 | 0.83498 | 0.00285 |
| CCL20/MIP-3 alpha | 15.5656698 | 13.0605 | 1.221683 | 0.30985 |
| CCL3/CCL4 MIP-1 alpha/beta | 19.7142228 | 15.0717 | 1.440288 | 0.263417 |
| CCL5/RANTES | 17.3851902 | 5.397992 | 1.390715 | 0.193217 |
| CCL7/MCP-3 | 4.6860314 | 3.252532 | 0.838003 | 0.068895 |
| CD14 | 1.6636062 | 0.232924 | 1.109644 | 0.27837 |
| CD30 | 1.333497 | 0.169473 | 1.064739 | 0.049854 |
| CD31 | 1.3960622 | 0.161299 | 1.338063 | 0.485993 |
| CD40 ligand | 1.2925922 | 0.121816 | 1.081436 | 0.077479 |
| Chitinase 3-like 1 | 4.1547926 | 2.465684 | 0.702872 | 0.050424 |
| Complement Component C5/C5a | 1.6071846 | 0.23033 | 1.008338 | 0.154563 |
| Complement Factor D | 1.273009 | 0.065638 | 0.899029 | 0.056481 |
| Cripto-1 | 1.6708122 | 0.298568 | 1.015132 | 0.080083 |
| CXCL1/GRO alpha | 6.3920874 | 1.860376 | 0.978544 | 0.035433 |
| CXCL10/IP-10 | 89.180002 | 32.68489 | 1.218749 | 0.275235 |
| CXCL11/I-TAC | 41.7030432 | 33.17172 | 1.244408 | 0.385624 |
| CXCL12/SDF-1 alpha | 1.7287528 | 0.181067 | 1.040522 | 0.179947 |
| CXCL4/PF4 | 1.605583 | 0.178836 | 1.260547 | 0.250934 |
| CXCL5/ENA-78 | 13.9085766 | 9.885841 | 0.944036 | 0.023785 |
| CXCL9/MIG | 2.6364888 | 0.942528 | 1.396962 | 0.168094 |
| Cystatin C | 2.3061618 | 0.506042 | 1.151898 | 0.134713 |
| Dkk-1 | 4.965894 | 1.474081 | 1.089239 | 0.057725 |
| DPPIV/CD26 | 1.7739458 | 0.320999 | 1.021454 | 0.076379 |
| EGF | 2.1586222 | 0.713946 | 1.01189 | 0.062607 |
| EMMPRIN | 2.5718048 | 0.789068 | 1.123027 | 0.108206 |
| Endoglin/CD105 | 1.5065322 | 0.203504 | 0.762404 | 0.024847 |
| Fas Ligand | 1.3421876 | 0.165382 | 0.90571 | 0.049545 |
| FGF basic | 1.4966196 | 0.105137 | 0.937225 | 0.065459 |
| FGF-19 | 1.657135 | 0.210628 | 1.190624 | 0.115651 |

| | | | | |
|---|---|---|---|---|
| Flt-3 Ligand | 1.8936294 | 0.353278 | 1.140806 | 0.040755 |
| G-CSF | 9.8155484 | 8.30316 | 1.128102 | 0.041315 |
| GDF-15 | 4.8741136 | 1.763071 | 0.948506 | 0.068209 |
| GM-CSF | 1.6934694 | 0.378477 | 1.126109 | 0.061572 |
| Growth Hormone (GH) | 1.7356002 | 0.274018 | 0.810799 | 0.185592 |
| HGF | 4.2018714 | 2.007048 | 0.881435 | 0.128722 |
| ICAM-1/CD54 | 2.1747818 | 0.557003 | 0.950961 | 0.0408 |
| IFN-gamma | 1.2768538 | 0.054068 | 0.970812 | 0.09473 |
| IGFBP-2 | 1.377401 | 0.14277 | 1.040726 | 0.026047 |
| IGFBP-3 | 4.034496 | 2.15735 | 1.102966 | 0.019377 |
| IL-1 alpha/IL-1F1 | 1.2858292 | 0.088766 | 1.228784 | 0.106472 |
| IL-1 beta/IL-1F2 | 1.7747486 | 0.362789 | 1.131139 | 0.035275 |
| IL-10 | 1.5192256 | 0.099035 | 0.969178 | 0.086008 |
| IL-11 | 1.4471918 | 0.095497 | 1.009244 | 0.100844 |
| IL-12 p70 | 1.3750594 | 0.10219 | 1.093594 | 0.085481 |
| IL-13 | 1.696196 | 0.339707 | 1.206015 | 0.103586 |
| IL-15 | 1.9582918 | 0.460879 | 1.08062 | 0.079593 |
| IL-16 | 2.5615664 | 0.635304 | 1.087178 | 0.140808 |
| IL-17A | 1.9654252 | 0.418088 | 1.126834 | 0.163742 |
| IL-18 BPa | 15.7316144 | 13.57683 | 1.305137 | 0.355774 |
| IL-19 | 1.4159508 | 0.169109 | 0.980446 | 0.016779 |
| IL-1ra/IL-1F3 | 1.9606684 | 0.439469 | 1.048213 | 0.092304 |
| IL-2 | 1.5766846 | 0.393835 | 1.141399 | 0.079841 |
| IL-22 | 1.4119306 | 0.125007 | 0.782652 | 0.15752 |
| IL-23 | 1.4587832 | 0.193636 | 0.934651 | 0.101298 |
| IL-24 | 1.5186352 | 0.104092 | 0.97378 | 0.077665 |
| IL-27 | 1.4768848 | 0.117661 | 1.159607 | 0.262546 |
| IL-3 | 1.4575902 | 0.34632 | 1.139685 | 0.147909 |
| IL-31 | 1.5169752 | 0.212865 | 1.060714 | 0.190246 |
| IL-32 alpha/beta/gamma | 1.4680872 | 0.154683 | 1.152776 | 0.121204 |
| IL-33 | 1.665933 | 0.329233 | 1.233423 | 0.003236 |
| IL-34 | 2.3314664 | 0.545274 | 1.235241 | 0.091049 |
| IL-4 | 1.5043808 | 0.171366 | 0.997863 | 0.026452 |
| IL-5 | 1.2458654 | 0.137511 | 0.799691 | 0.180663 |
| IL-6 | 10.3800752 | 7.209792 | 0.93069 | 0.099206 |
| IL-8 | 11.86945 | 4.245107 | 0.917451 | 0.036495 |
| Kallikrein 3/PSA | 1.679024 | 0.289404 | 1.422937 | 0.528298 |
| KGF/FGF-7 | 1.6478468 | 0.209421 | 0.997695 | 0.044396 |
| Leptin | 1.4187688 | 0.148598 | 0.960115 | 0.013242 |
| LIF | 1.2874882 | 0.152914 | 0.806076 | 0.199003 |
| Lipocalin-2/NGAL | 1.396939 | 0.110544 | 0.929148 | 0.108902 |

| | | | | |
|---|---|---|---|---|
| M-CSF | 1.5989998 | 0.212569 | 0.996407 | 0.158766 |
| MIF | 1.8563596 | 0.610488 | 1.205771 | 0.036568 |
| MMP-9 | 6.5277048 | 2.745464 | 1.041292 | 0.289401 |
| Myeloperoxidase | 1.2290274 | 0.142744 | 0.938321 | 0.153145 |
| Osteopontin (OPN) | 2.9220832 | 1.044759 | 0.849462 | 0.228904 |
| PDGF-AA | 1.6000452 | 0.445003 | 0.899309 | 0.132587 |
| PDGF-AB/BB | 1.500799 | 0.074826 | 0.997536 | 0.155006 |
| Pentraxin-3 | 2.5716456 | 1.054416 | 0.839467 | 0.027348 |
| RAGE | 1.8837892 | 0.363125 | 2.145745 | 1.015983 |
| RBP4 | 2.0805912 | 0.439306 | 1.240603 | 0.149995 |
| Relaxin-2 | 1.7922062 | 0.212301 | 1.415175 | 0.501026 |
| Resistin | 1.6785924 | 0.196995 | 1.021107 | 0.212882 |
| Serpin E1/PAI-1 | 2.5615458 | 0.902751 | 0.797816 | 0.073465 |
| SHBG | 1.4661384 | 0.102449 | 1.112722 | 0.002808 |
| ST2/IL-1 R4 | 1.2581856 | 0.073922 | 0.930787 | 0.23042 |
| TFF3 | 1.4036804 | 0.058148 | 1.069102 | 0.279255 |
| TfR | 1.570087 | 0.278103 | 1.417126 | 0.55259 |
| TGF-alpha | 1.6747386 | 0.322899 | 1.630656 | 0.478111 |
| Thrombospondin-1 | 1.4619158 | 0.26512 | 1.184769 | 0.051603 |
| TIM-3 | 1.4701726 | 0.270349 | 1.65791 | 0.882127 |
| TNF-alpha | 1.5956582 | 0.279569 | 1.212706 | 0.101393 |
| uPAR | 2.348538 | 0.402858 | 1.365572 | 0.466394 |
| VCAM-1 | 1.7173468 | 0.440618 | 1.103584 | 0.246384 |
| VEGF | 7.2772 | 2.599381 | 1.358042 | 0.576299 |
| Vitamin D BP | 1.211376 | 0.113223 | 1.188577 | 0.223708 |
