## Supplemental Table 4 for "Innate Immune Activation and Mitochondrial ROS Invoke Persistent Cardiac Conduction System Dysfunction after COVID-19"

### Supplementary Table 4: additional statistical tests

Figure2

Adj. p-values

| | | DPI | D1 | D3 | D5 | D7 | D14 | D21 | D28 |
|---|---|---|---|---|---|---|---|---|---|
| 2B | SARS-CoV2 | compared to baseline (D0) | <0.0001 | <0.0001 | <0.0001 | 0.7875 | 0.01409 | 0.0013 | <0.0001 |
| | | compared to Mock | <0.0001 | <0.0001 | <0.0001 | 1 | 0.9995 | 0.0254 | 0.0019 |
| 2C | SARS-CoV2 | compared to baseline | 0.0014 | 1 | 0.0028 | 0.0028 | 0.4802 | 0.1841 | 0.0266 |
| | | compared to Mock | 0.0005 | | 0.0015 | 0.0005 | | 0.012 | 0.001 |
| 2D | SARS-CoV2 | compared to baseline | <0.0001 | 0.0021 | 0.2917 | 0.014 | 0.3395 | 0.0168 | <0.0001 |
| | | compared to Mock | 0.001 | 0.01 | | | | 0.0095 | <0.0001 |
| 2E | SARS-CoV2 | compared to baseline | <0.0001 | <0.0001 | <0.0001 | <0.0001 | 0.0014 | <0.0001 | 0.0007 |
| | | compared to Mock | <0.0001 | <0.0001 | <0.0001 | <0.0001 | <0.0001 | 0.004 | 0.00125 |

Figure 7

| | | DPI | 1 | 2 | 3 | 4 | 5 | 6 | 7 | 8 |
|---|---|---|---|---|---|---|---|---|---|---|
| 7A Body Weight | SARS-CoV2 | compared to baseline | 1 | 0.8325 | 0.0005 | <.0001 | <.0001 | <.0001 | <.0001 | <.0001 |
| | | compared to Mock | 1 | 0.9577 | 0.0004 | <.0001 | <.0001 | <.0001 | <.0001 | <.0001 |
| | mTEMPO | compared to baseline | 1 | 0.9998 | 0.212 | 0.0045 | 0.0003 | <.0001 | 0.0003 | 0.0466 |
| | | compared to Mock | 1 | 0.9996 | 0.0302 | <.0001 | <.0001 | <.0001 | <.0001 | <.0001 |
| | | compared to SARS-CoV2 | 1 | 1 | 1 | 1 | 1 | 1 | 1 | 1 |
| | Ruxo | compared to baseline | 1 | 0.9974 | 0.0075 | <.0001 | <.0001 | <.0001 | <.0001 | <.0001 |
| | | compared to Mock | 1 | 0.9972 | 0.0006 | <.0001 | <.0001 | <.0001 | <.0001 | <.0001 |
| | | compared to SARS-CoV2 | 1 | 1 | 1 | 1 | 1 | 1 | 1 | 1 |

| | | DPI | 9 | 10 | 11 | 12 | 13 | 14 | 21 | 28 |
|---|---|---|---|---|---|---|---|---|---|---|
| | SARS-CoV2 | compared to baseline | 0.0007 | 0.0285 | 0.9022 | 1 | 1 | . | 0.2229 | <.0001 |
| | | compared to Mock | <.0001 | <.0001 | <.0001 | 0.0003 | 0.003 | . | 0.063 | 0.9839 |
| | mTEMPO | compared to baseline | 0.7285 | 0.9914 | 1 | 1 | 1 | 1 | 0.2465 | 0.0008 |
| | | compared to Mock | <.0001 | 0.0004 | 0.0033 | 0.0188 | 0.1054 | 0.7395 | 0.99 | 0.6869 |
| | | compared to SARS-CoV2 | 1 | 1 | 1 | 1 | 1 | . | 1 | 1 |
| | Ruxo | compared to baseline | 0.0002 | 0.022 | 0.9797 | 1 | 1 | 1 | 0.0917 | <.0001 |
| | | compared to Mock | <.0001 | <.0001 | <.0001 | 0.0013 | 0.0085 | 0.7219 | 0.9834 | 1 |
| | | compared to SARS-CoV2 | 1 | 1 | 1 | 1 | 1 | . | 1 | 1 |

| | | DPI | D1 | D3 | D5 | D7 | D14 | D21 | D28 |
|---|---|---|---|---|---|---|---|---|---|
| 7B Body Temperature | SARS-CoV2 | compared to baseline | 0.7425 | <.0001 | 0.0003 | 0.8342 | 0.9839 | 0.4971 | 0.8977 |
| | | compared to Mock | 0.767 | 0.0002 | 0.0069 | 0.9997 | 1 | 0.9905 | 0.9907 |
| | mTEMPO | compared to baseline | 0.915 | 0.0039 | 0.5111 | 0.9978 | 0.9984 | 0.986 | 0.9898 |
| | | compared to Mock | 0.8985 | 0.0148 | 0.7792 | 1 | 1 | 1 | 0.9993 |
| | | compared to SARS-CoV2 | 1 | 1 | 1 | 1 | 1 | 1 | 1 |
| | Ruxo | compared to baseline | 0.148 | <.0001 | 0.0002 | 0.9971 | 1 | 1 | 1 |
| | | compared to Mock | 0.0212 | <.0001 | <.0001 | 0.9994 | 1 | 1 | 1 |
| | | compared to SARS-CoV2 | 0.6568 | 0.9604 | 0.7034 | 1 | 1 | 1 | 1 |

Figure 7 (continued)

| | | DPI | D1 | D3 | D5 | D7 | D14 | D21 | D28 |
|---|---|---|---|---|---|---|---|---|---|
| 7C Breathing Rate | SARS-CoV2 | compared to baseline | 1 | <.0001 | <.0001 | <.0001 | 0.9958 | 1 | 0.9999 |
| | | compared to Mock | 0.7157 | <.0001 | <.0001 | <.0001 | 0.1008 | 1 | 1 |
| | mTEMPO | compared to baseline | 1 | <.0001 | <.0001 | <.0001 | 0.988 | 1 | 1 |
| | | compared to Mock | 1 | <.0001 | <.0001 | <.0001 | 0.3383 | 0.9992 | 1 |
| | | compared to SARS-CoV2 | 0.9969 | 0.0003 | <.0001 | <.0001 | 1 | 1 | 1 |
| | Ruxo | compared to baseline | 1 | <.0001 | <.0001 | <.0001 | <.0001 | 1 | 1 |
| | | compared to Mock | 1 | <.0001 | <.0001 | <.0001 | <.0001 | 1 | 1 |
| | | compared to SARS-CoV2 | 0.9919 | 0.0034 | <.0001 | <.0001 | 0.7959 | 1 | 1 |
| 7D RR Interval | mTEMPO | compared to baseline | <0.0001 | <0.0001 | <0.0001 | 0.8807 | 0.0784 | 0.0007 | 0.0002 |
| | | compared to Mock | <0.0001 | <0.0001 | <0.0001 | 1 | 0.9997 | 0.1726 | 0.185 |
| | | compared to SARS-CoV2 | <0.0001 | 0.0433 | 1 | 1 | 1 | 1 | 1 |
| | Ruxo | compared to baseline | <0.0001 | <0.0001 | <0.0001 | 0.2308 | 0.2083 | 0.0014 | <0.0001 |
| | | compared to Mock | <0.0001 | <0.0001 | <0.0001 | 1 | 1 | 0.3091 | 0.0283 |
| | | compared to SARS-CoV2 | 1 | 1 | 0.6173 | 1 | 1 | 1 | 1 |
| 7E RR100 | mTEMPO | compared to baseline | 0.021 | 0.2842 | 0.7798 | 0.161 | 1 | 1 | 0.5873 |
| | | compared to Mock | 0.00567 | | 0.04983 | 0.00433 | | 0.178 | 0.0935 |
| | | compared to SARS-CoV2 | 0.15275 | | 0.0536 | 0.1275 | | 0.07033 | 0.025 |
| | Ruxo | compared to baseline | <0.0001 | 0.0301 | 0.0301 | 0.0301 | 0.0798 | 0.2009 | 0.0112 |
| | | compared to Mock | 0.00175 | | 0.0055 | 0.00433 | | 0.4391 | 0.0555 |
| | | compared to SARS-CoV2 | 0.4745 | | 0.429 | 0.222 | | 0.028 | 0.0555 |
| 7F RMSSD | mTEMPO | compared to baseline | 0.0372 | 0.1673 | 1 | 0.8953 | 1 | 0.3584 | 0.2982 |
| | | compared to Mock | 0.005 | 0.0103 | | | | 0.0095 | 0.02525 |
| | | compared to SARS-CoV2 | 0.3368 | 0.3871 | | | | 0.3147 | 0.02525 |
| | Ruxo | compared to baseline | 0.0354 | 0.1876 | 1 | 0.3472 | 1 | 1 | 0.7035 |
| | | compared to Mock | 0.001 | 0.01 | | | | 0.0095 | 0.00025 |
| | | compared to SARS-CoV2 | 0.227375 | 0.202 | | | | 0.3147 | 0.2499 |
| 7G AV block | mTEMPO | compared to baseline | <0.0001 | <0.0001 | <0.0001 | 0.0112 | 0.0735 | 0.0742 | 0.7294 |
| | | compared to Mock | 0.00517 | 0.002 | 0.00283 | 0.032 | 0.1035 | 0.305875 | 0.4939 |
| | | compared to SARS-CoV2 | 0.0261 | 0.00225 | 0.011875 | 0.008 | 0.004833 | 0.01667 | 0.002167 |
| | Ruxo | compared to baseline | <0.0001 | 0.0021 | 0.0091 | 0.4501 | 0.29246 | 1 | 0.3248 |
| | | compared to Mock | 0.00175 | 0.002 | 0.00283 | 0.02017 | 0.132 | 0.5 | 0.285375 |
| | | compared to SARS-CoV2 | 0.1656 | 0.0087 | 0.0279 | 0.031 | 0.004833 | 0.01025 | 0.00125 |
